# Graph of graphs analysis for multiplexed data with application to imaging mass cytometry

**DOI:** 10.1101/2020.08.23.263467

**Authors:** Ya-Wei Eileen Lin, Tal Shnitzer, Ronen Talmon, Franz Villarroel-Espindola, Shruti Desai, Kurt Schalper, Yuval Kluger

## Abstract

Hyper spectral imaging, sensor networks, spatial multiplexed proteomics, and spatial transcriptomics assays is a representative subset of distinct technologies from diverse domains of science and engineering that share common data structures. The data in all these modalities consist of high-dimensional multivariate observations (*m*-dimensional feature space) collected at different spatial positions and therefore can be analyzed using similar computational methodologies. Furthermore, in many studies practitioners collect datasets consisting of multiple spatial assays of this type, each capturing such data from a single biological sample, patient, or hyper spectral image, etc. Each of these spatial assays could be characterized by several regions of interest (ROIs). The focus of this paper is on a particular application, imaging mass cytometry (IMC), which falls into this problem setup. To extract meaningful information from the multi-dimensional observations recorded at different ROIs across different assays, we propose to analyze such datasets using a two-step graph-based approach. We first construct for each ROI a graph representing the interactions between the *m* covariates and compute an *m* dimensional vector characterizing the steady state distribution among features. We then use all these *m*-dimensional vectors to construct a graph between the ROIs from all assays. This second graph is subjected to a nonlinear dimension reduction analysis, retrieving the intrinsic geometric representation of the ROIs. Such a representation provides the foundation for efficient and accurate organization of the different ROIs that correlates with their phenotypes. Theoretically, we show that when the ROIs have a particular bi-modal distribution, the new representation gives rise to a better distinction between the two modalities compared to the maximum a posteriori (MAP) estimator. We applied our method to predict the sensitivity to PD-1 axis blockers treatment of lung cancer subjects based on IMC data, achieving 92% accuracy. This serves as empirical evidence that the graph of graphs approach enables us to integrate multiple ROIs and the intra-relationships between the features at each ROI, giving rise to an informative representation that is strongly associated with the phenotypic state of the entire image. Importantly, this approach is applicable to other modalities such as spatial transcriptomics.

**Author summary:** We propose a two-step graph-based analyses for high-dimensional multiplexed datasets characterizing ROIs and their inter-relationships. The first step consists of extracting the steady state distribution of the random walk on the graph, which captures the mutual relations between the covariates of each ROI. The second step employs a nonlinear dimensionality reduction on the steady state distributions to construct a map that unravels the intrinsic geometric structure of the ROIs. We show theoretically that when the ROIs have a two-class structure, our method accentuates the distinction between the classes. Particularly, in a setting with Gaussian distribution it outperforms the MAP estimator, implying that the mutual relations between the covariates and spatial coordinates are well captured by the steady state distributions. We apply our method to imaging mass cytometry (IMC). Our analysis provides a representation that facilitates prediction of the sensitivity to PD-1 axis blockers treatment of lung cancer subjects. Particularly, our approach achieves state of the art results with accuracy of 92%.

## Introduction

Consider multi-feature observations collected at different spatial positions. Data structure of this type requires analysts to address two immediate natural questions. First is how to characterize the associations between the different features in each position. Second is how to organize the observations from different spatial positions into an informative representation.

We approach these two questions from the standpoint of manifold learning, which is a class of nonlinear dimensionality reduction techniques for high-dimensional data [38, 32, 3, 10]. The common assumption in manifold learning is that the multi-feature observations lie on a hidden lower-dimensional manifold. Such an assumption facilitates the incorporation of geometric concepts such as metrics, geodesic distances, and embedding, into useful data analysis techniques. In order to learn a (continuous) manifold from discrete data samples, commonly-used manifold learning methods rely on the construction of a graph. Typically, the data samples form the graph nodes and the edges of the graph are determined according to some similarity notion that is usually application-specific.

In our work, we adhere to manifold learning techniques and propose a method consisting of two-step graph analysis. At the first stage, we build a graph for each spatial position, where the graph nodes are the multi-feature observations. The motivation to build such a graph rather than using the observations directly stems from an assumption that the information about the sample at each spatial position is better expressed by the mutual-relations between the features. Then, we define a random walk on this graph and build a characteristic vector of the respective spatial position by computing the steady state distribution (SSD) of the random walk. For analysis purposes, we define a new notion of heterogeneity, representing a statistical diversity of the multiple features, and show that the SSD characterises each spatial position in terms of this heterogeneity. In addition, using this notion of heterogeneity, when the density of the observations at the spatial positions is bi-modal, we show that these SSDs can lead to an accurate identification of the two modes, outperforming the maximum a-posteriori (MAP) estimator [28] in a statistical setting with Gaussian distributions.

At the second stage, we build a graph whose nodes are the new characteristic vectors (SSDs) of all the spatial positions. We apply diffusion maps [10] to this second graph and obtain a low dimensional representation of the spatial positions. The dimension of the computed representation is determined by a nonlinear variant of the Jackstraw algorithm [9].

Broadly, the proposed algorithm could be viewed as building a *graph of graphs*. From a manifold learning standpoint, this two-step procedure could be viewed as inferring a *manifold of manifolds*. Namely, at the first stage, we recover the local manifolds that underlie the multiple features at each spatial location, and then, at the second stage, we recover the global manifold between the spatial positions, formed by the collection of all local manifolds. This standpoint is related to a large body of recent work involving the discovery and analysis of multi-manifold structures, e.g., alternating diffusion [20, 37, 34, 16], multi-view diffusion maps [21], joint Laplacian diagonalization [14], to name just a few. Therefore, the proposed method can be viewed as a follow up work along this line of research.

We apply our method to imaging mass cytometry (IMC) [5, 15, 7]. IMC is a new technique for multiplexed simultaneous imaging of proteins and protein modifications at subcellular resolution aimed for uncovering pathologies in tissues and tumors. IMC analysis can also be used to study the composition of non-diseases tissue samples such as histology studies or molecular profiles. The acquired intensities of the protein expression levels are viewed as markers, providing important biological information on the tissues of interest. This acquisition procedure gives rise to multi-feature observations at different spatial positions, where the multiple features are the markers and the spatial positions are ROIs within pathology slides. Our experimental study focuses on one of the important tasks of IMC data analysis: associating the response status of a patient to a therapeutic intervention with a high-dimensional spatial IMC sample from the relevant patients’ tissues. Here, we propose to recast this problem as a binary hypothesis testing problem. We assume that all the ROIs of each patient can be labeled by the patient’s response or non-response status. The collection of all ROIs from the patients’ cohort induces a bi-modal density of expression signatures. Then, given the spatial protein expression levels of a certain tissue type, we ask whether the subject was responsive to treatment. We showcase the performance of the proposed method on two IMC cohorts consisting of samples taken from lung cancer subjects. We achieve a 92% prediction accuracy of response to treatment (PD-1 axis blockers) in an unsupervised manner. This result outperforms competing methods, specifically, the results obtained by diffusion maps [10] and by the well-known heat kernel signature (HKS) [35].

## Results

### Overview

We start by presenting the problem setting. Consider *n* data points 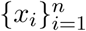 from a hidden manifold ℳ embedded in a high-dimensional Euclidean space ℝ^*k*^. Assume we do not have direct access to these data points; instead, these data points are measured through *m* observation functions *f*_*j*_ : ℳ → ℝ, where *j* = 1, …, *m* is the index of the observation function. Given *n* multi-feature observations *f*_*j*_(*x*_*i*_) of the data point *x*_*i*_ for *i* = 1, …, *n*, each consisting of *m* features *j* = 1, …, *m*, our goal is to recover the data points *x*_*i*_ on the hidden manifold ℳ.

In the context of IMC, the data points represent the treatment outcome based on *n* spatial positions located at ROIs within pathology slides of tissues from several patients. At each spatial position *i* = 1, …, *n*, the observations *f*_*j*_(*x*_*i*_) for *j* = 1, …, *m* are the expression levels of *m* markers. Each observation *f*_*j*_(*x*_*i*_) ∈ℝ^*d*^ is a patch of *d* pixels of the expression level image of marker *j* at position *i*.

To simplify the presentation of our approach, we begin with an illustrative localization problem, which is simpler than the IMC problem. Suppose we have a surface ℳ and objects located at *x*_*i*_ on ℳ. The locations *x*_*i*_ are hidden, but measured through *m* sensors, such that for each location *x*_*i*_ we have *m* multi-feature observations 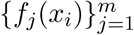. That is *f*_*j*_(*x*_*i*_) is a *d*-dimensional observation of sensor *j* when the object is at *x*_*i*_. The goal is to recover the locations *x*_*i*_ on the surface ℳ given 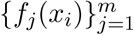.

Our approach consists of two stages. At the first stage, we construct a graph for each data point *x*_*i*_ in order to capture associations between its *m* multi-feature observations 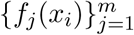. Capturing such mutual-relationships is natural in the context of localization problems, e.g., the triangulation property [19] in which the relative locations of the sensors are exploited. In addition, these mutual-relations are typically more robust to noise in comparison with the nominal values of the multi-feature observations, *f*_*j*_(*x*_*i*_), themselves. Concretely, consider the *m* observations 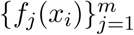 associated with the data point *x*_*i*_. Each observation *f*_*j*_(*x*_*i*_) forms a single node in the graph, hereby denoted as node *j*, giving rise to a graph with a total of *m* nodes. The graph we consider is the complete graph, where the weights of the edges are determined based on the Euclidean distance between the corresponding observations: the weight of the edge connecting nodes *j* and *k* is proportional to exp{−∥*f*_*j*_(*x*_*i*_) − *f*_*k*_(*x*_*i*_) ∥^2^} Then, we compute the SSD of a random walk defined on this graph at each location. SSD has a vector form that embodies the multi-feature inter-relationships of the data point *x*_*i*_.

At the next step, we define a second graph based on the SSDs, characterizing the points 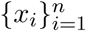. Concretely, each data point *x*_*i*_ is represented by a node, and the pairwise distances between the SSDs form the adjacency matrix of the graph. Then, we apply a particular manifold learning technique, diffusion maps [10], to this graph. This application facilitates the recovery of the hidden manifold ℳ in the sense that an embedding of the points *x*_*i*_ is learned, such that the distances between the embedded points respect a notion of an intrinsic distance (the diffusion distance [10]) on ℳ. The application of diffusion maps to the second graph in the context of the localization problem gives rise to an embedding that serves as an accurate representation of the hidden locations of the data point. In Section Localization toy problem, we demonstrate the proposed method on several simulations of such a localization toy problem.

We remark that the IMC problem and the localization problem share many aspects. For example, in both problems, the multi-feature observations are noisy and the mutual-relationship between them carry important information. Yet, there is a particular aspect that makes the IMC problem more challenging; while the points on the hidden manifold in a localization problem are homogeneous because they all represent location coordinates, the points in the IMC problem could be significantly different due to the large variability in the tissue structure. Importantly, the proposed method accommodates the joint processing of such different points through their representation by the SSD.

### Proposed Method

The first step of the proposed method is to construct an undirected weighted graph *𝒢*_*i*_ = (*𝒱*_*i*_, *ε*_*i*_, ***W***_*i*_) for each data point *x*_*i*_ ∈ ℳ ⊂ ℝ^*k*^, where the vertex set is *𝒱*_*i*_ = {*f*_1_(*x*_*i*_), *f*_2_(*x*_*i*_), *…, f*_*m*_(*x*_*i*_)*}*, the edge set is *ε*_*i*_ *⊆𝒱*_*i*_ × *𝒱*_*i*_, and the graph weights matrix ***W***_*i*_ ∈ℝ^*m*×*m*^ is given by the Gaussian kernel

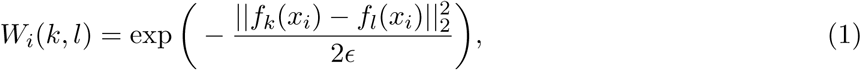

where *k, l* ∈ {1, *…, m}* and *ϵ >* 0 is a scale parameter. Note that since ***W***_*i*_ is symmetric, ***W***_*i*_ is diagonalizable. That is, there is a set of real eigenvalues 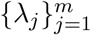 with a corresponding orthonormal basis of eigenvectors 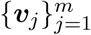 such that

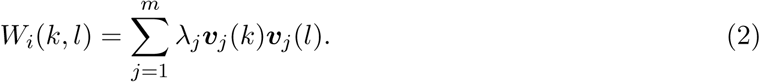

Next, we define a random walk on the graph *𝒢*_*i*_. Let ***P***_*i*_ ∈ℝ^*m*×*m*^ be a row stochastic matrix given by

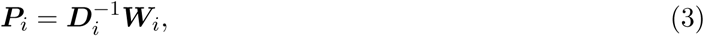

where ***D***_*i*_ is a diagonal matrix whose diagonal elements are given by 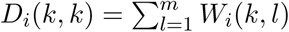. The value of *P*_*i*_(*k, l*) can be interpreted as a transition probability from a vertex *f*_*k*_(*x*_*i*_) to a vertex *f*_*l*_(*x*_*i*_) in one step of a random walk on the graph *𝒢*_*i*_.

The transition probability matrix ***P***_*i*_ is self-adjoint and compact, and therefore, the spectral decomposition of 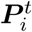 for *t >* 0 is given by

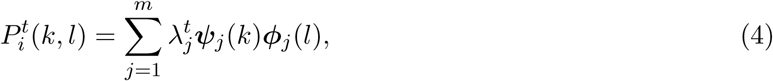

where 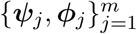 are the right-and left-eigenvectors with the corresponding eigenvalues 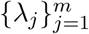. By the construction of ***P***_*i*_ from ***W***_*i*_, the relations between their respective eigenvectors are given by

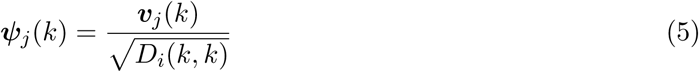

and

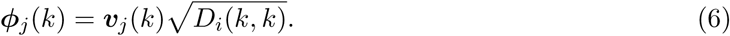

Interestingly, in the special case where *t* = 1, the probability of the node *f*_*k*_(*x*_*i*_) to stay in place is given by

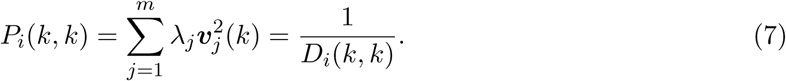

Note that ***ϕ***_1_ ∈ℝ^*m*^ is the left eigenvector of ***P***_*i*_ corresponding to *λ*_1_ = 1, satisfying:

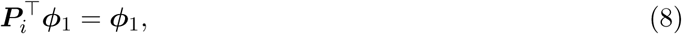

where 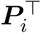 is the transpose of the matrix ***P***_*i*_. Consider an arbitrary distribution vector ***π***_0_ ∈ℝ^*m*^ and observe the expansion:

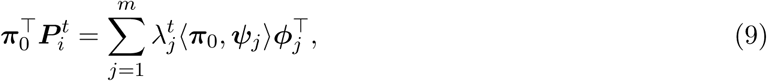

where 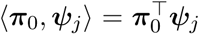 is the standard Euclidean product. In the limit 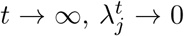 for *j* > 1 and therefore

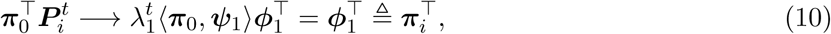

where ***π***_*i*_ ∈ℝ^*m*^since *λ*_1_ = 1, ***ψ***_1_ = **1**, and 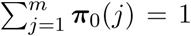. Since the random walk defined by ***P***_*i*_ is irreducible, finite, and aperiodic [23], the stationary distribution ***π***_*i*_ is a *unique* stationary distribution. The convergence in Eq (10) and the uniqueness allow us to treat the stationary distribution in this case as the steady state distribution (SSD). Note that the SSD ***π***_*i*_ *∝* ***D***_*i*_**1**, where **1** ∈ℝ^*m*^ is an all-ones vector. In other words, ***π***_*i*_ can be viewed as a normalized degrees vector of the graph *𝒢*_*i*_.

We will use ***π***_*i*_ as a new characteristic vector, or a signature, of *x*_*i*_, and consequently, the induced pairwise distances ∥***π***_*i*_ − ***π***_*i*′_ ∥, where *i, i*^*′*^ ∈ 1, …, *n*, will be used as the desired distances between the respective graphs *𝒢*_*i*_ and *𝒢*_*i*′_ for recovering ℳ. At first glance, using ***π***_*i*_ may seem too simplistic. Instead, one could use the broad spectral information. Consequently, define the Diffusion Kernel Signature (DKS) by

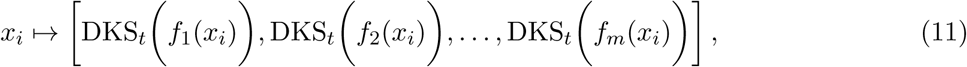

where

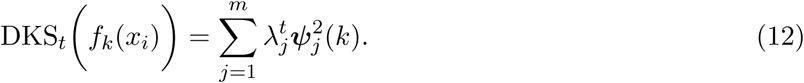

Since *λ*_*j*_ is in descending order and in [0, 1], the weight they assign to the eigenvectors in (12) becomes smaller as *t* increases. As a result, the DKS can be viewed as a low-pass filter, which controls the spectral bandwidth. In addition, the DKS can be recast in terms of the diffusion distance, a notion of distance induced by diffusion maps [10] that was shown useful in a broad range of applications, e.g., [36, 12, 27]. For more details see Diffusion maps. Specifically, when *t* = 1, we can show that

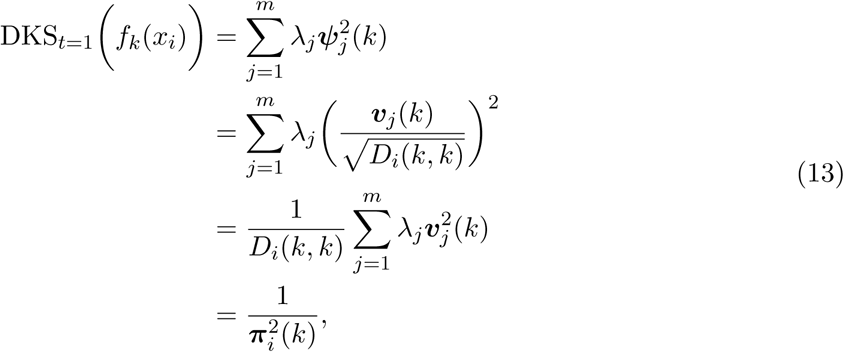

indicating that the SSD is a special case of DKS.

We note that DKS has already appeared in previous work in the context of spectral distances in [6, 8], where it was shown that it describes the underlying geometry of ℳ. We show in the following that the seemingly simple SSD, despite the lack of broad spectral information as in the DKS, still carries substantial information.

Note that *ϵ* is a scale parameter of the Gaussian kernel, where it can be used to infer locality. If *ϵ* is set to a small value, then ***π***_*i*_ captures local properties. Conversely, if *ϵ* is large, then ***π***_*i*_ represents the global structure. As a result, a multiscale signature can be formed, consisting of multiple SSDs *π*_*i*_ computed with different values of *ϵ*. An example is shown in S1 Fig.

The final stage of our method is building a low-dimensional representation of all the data points 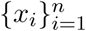. To this end, we apply diffusion maps to the corresponding characteristic vectors (signatures) 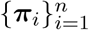 as follows. First, we build a global graph **G**^(2)^ whose nodes are ***π***_*i*_ and edge weights are determined by a Gaussian kernel based on the *l*_1_ *distance* between ***π***_*i*_.

That is, the global graph weights matrix **W**^(2)^ is defined by

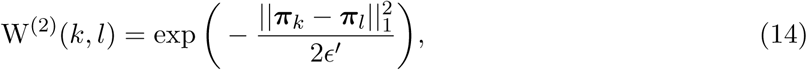

where *k, l* ∈ [1, *n*], ∥ · ∥_1_ is the *l*_1_ distance and *ϵ*^*′*^ *>* 0 is a scale parameter. We remark that common practice is to use the *l*_2_ distance in the Gaussian kernel. The reason we use the *l*_1_ distance is described in Binary Hypothesis Testing, which indeed leads to better empirical performance reported in Imaging mass cytometry (IMC).

Second, we construct a random walk, denoting its transition probability matrix by **P**^(2)^. Third, we apply the eigendecomposition to **P**^(2)^. Fourth, we set the dimension of the new representation according to the variant of the Jackstraw method [9], as shown in Determining the dimension of data.

The entire method is summarized in Algorithm 1 and a block diagram is illustrated in Fig 1.

**Figure 1:**
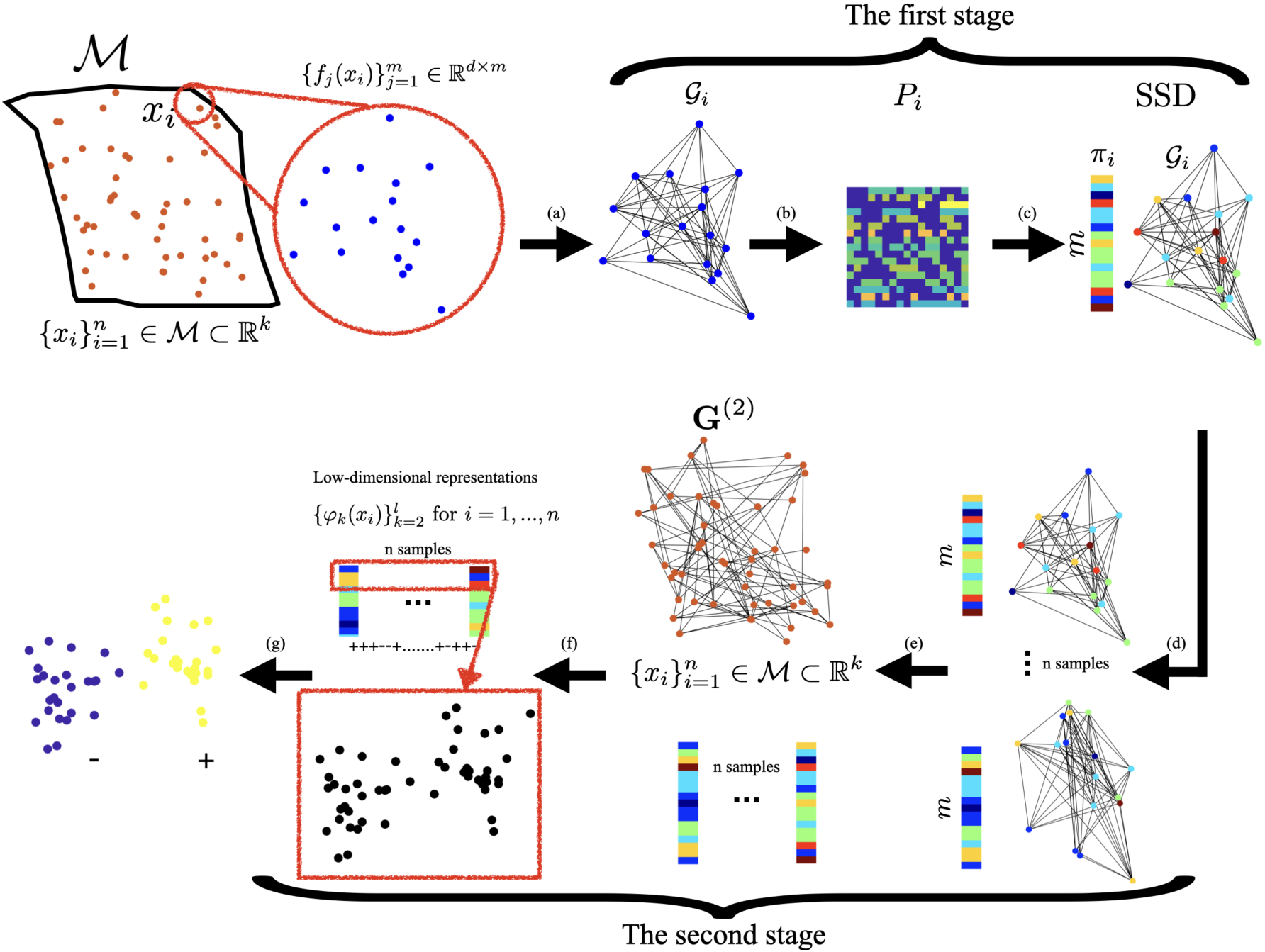
Illustrative diagram of Algorithm 1. (a) For each data point *x*_*i*_, we build a local graph *𝒢*_*i*_ based on its multi-feature observations 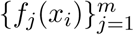. (b) We construct a random walk with transition probabilities matrix ***P***_*i*_ on *𝒢*_*i*_. (c) We extract the SSD signature ***π***_*i*_ from ***P***_*i*_. (e) We collect the SSDs of 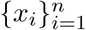 into an SSD representation matrix. (g) Subsequently, the matrix is subjected to a nonlinear dimensionality reduction using diffusion maps by the construction of the global graph **G**^(2)^ and the corresponding random walk with **P**^(2)^. (f) Via eigenvalue decomposition, we obtain a low-dimensional representation **Ψ**(*x*_*i*_) for *i* = 1, …, *n*, which is used in the subsequent tasks (g).

#### Algorithm 1

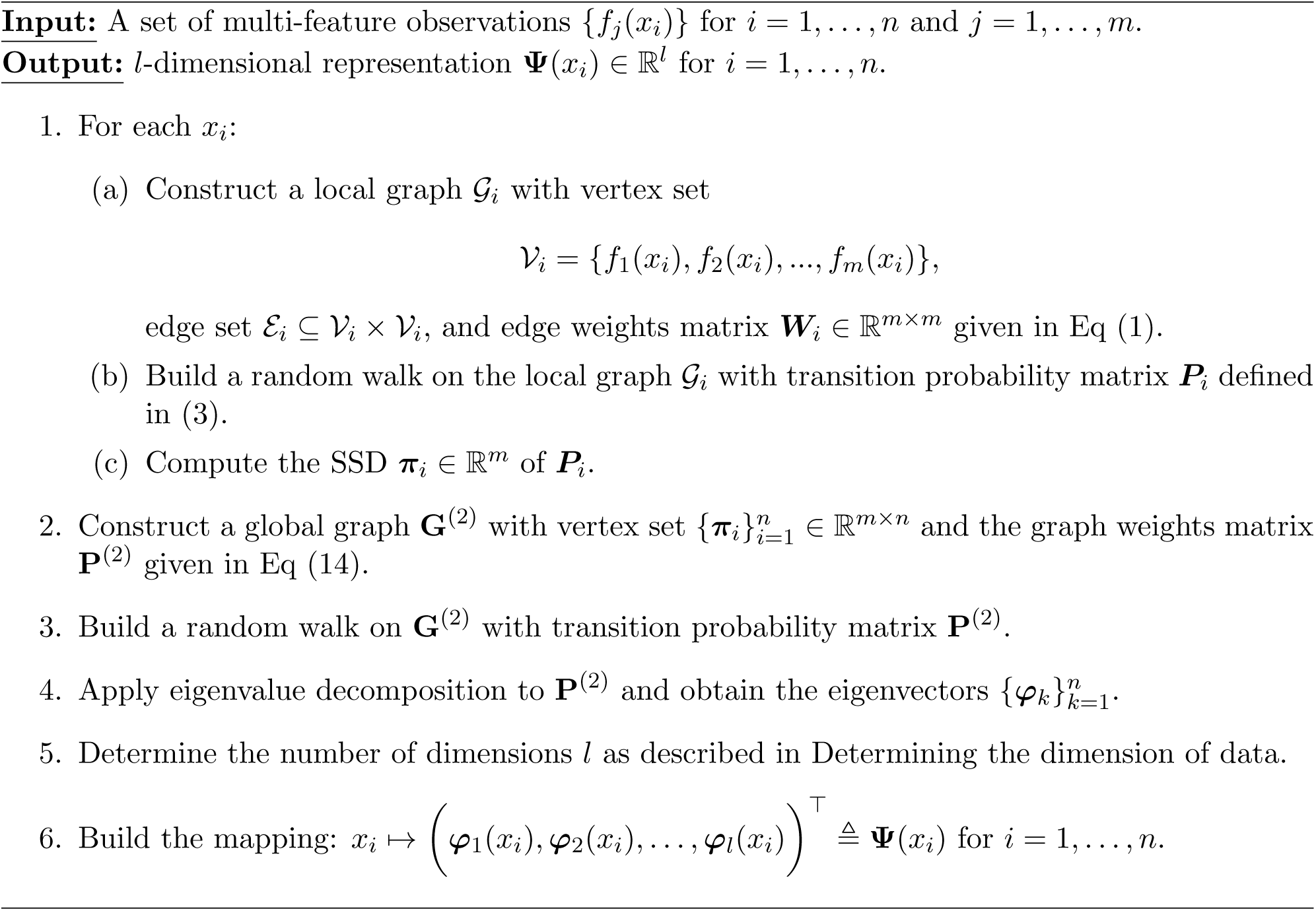

### Theoretical Analysis

We propose a statistical model that allows for a tractable analysis, showing the advantages of the SSD signature. Consider a data point *x*_*i*_ ∈ ℳ and denote the set of the observations by 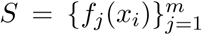. Assume the *j*-th observation *f*_*j*_(*x*_*i*_) ∈ℝ^*d*^ is a realization of a *d*-dimensional random vector 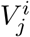 following a multivariate normal distribution given by

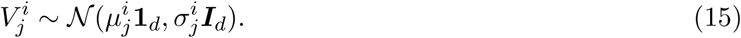

By collecting the *m* observations 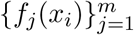 and denoting the covariance matrix between the *k* and *l* observation functions by 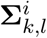, we obtain an *md*-dimensional vector that can be viewed as a realization of the random vector 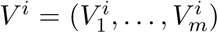 with the corresponding multivariate Gaussian distribution given by

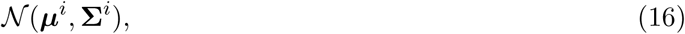

where

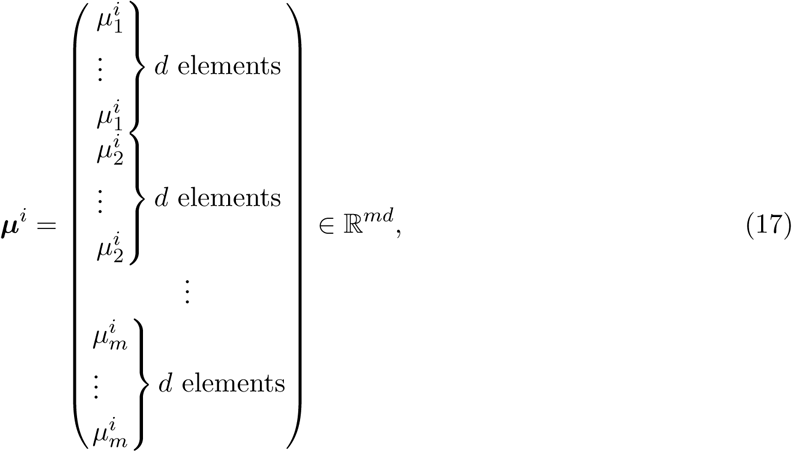

and

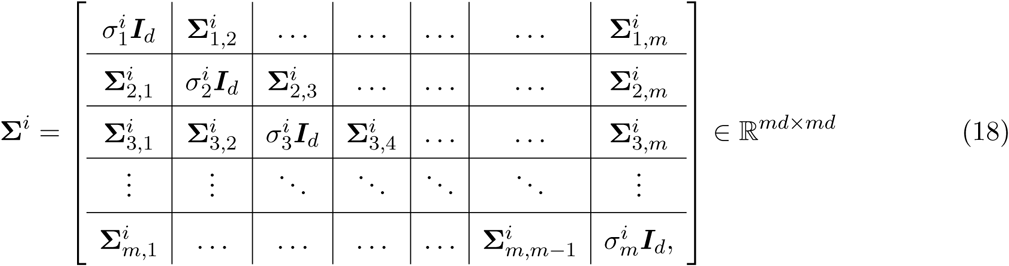

such that

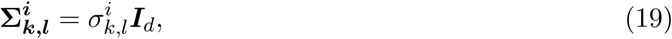

and 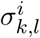 is the covariance between the *k*-th and *l*-th random vectors.

#### Definition 1

(empirical mean). *Given a set* Γ *and some real function on the set* ***q*** ∈ℝ^|Γ|^, *and a subset* Ω ⊂ Γ, *the empirical mean of* ***q*** *in* Ω *is defined by*

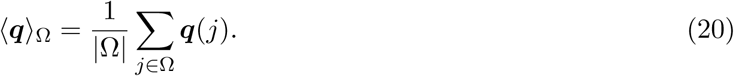

#### Definition 2

(heterogeneity). *Define the heterogeneity of a data point x*_*i*_ *by*

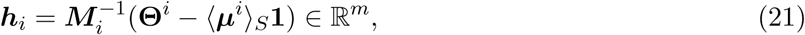

*where ⟨****μ*** ^*i*^*⟩* _*S*_ *is given by Definition 1*, ***M***_*i*_ *is an m* × *m diagonal matrix, whose diagonal elements are* 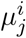, *and*

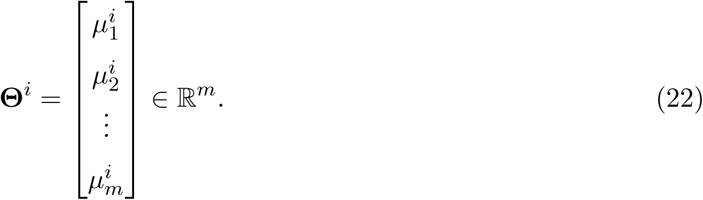

The heterogeneity ***h***_*i*_ ∈ℝ^*m*^ captures the mutual-relationships between the expected values of the observations 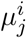 of a particular data point *x*_*i*_; if **Θ**(*j*)^*i*^ significantly deviates from *μ*^*i*^, then ***h***_*i*_(*j*) is large. Conversely, if **Θ**^*i*^(*j*) is close to *μ*^*i*^, then ***h***_*i*_(*j*) is close to zero.

#### Definition 3

(weighted heterogeneity). *Define* ***g***_*i*_ *as a weighted heterogeneity, whose jth element is given by*

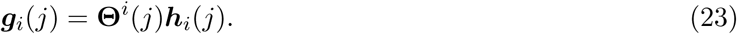

Under the considered statistical model, with the above definitions, the SSD ***π***_*i*_ can be written explicitly.

#### Proposition 1

*The j-th element of* ***π***_*i*_ *can be approximated by*

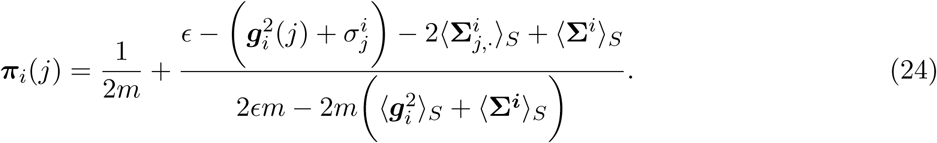

The derivation is based on the Taylor expansion of the Gaussian function in Eq (1), where *ϵ* is the scale of the function. The proof appears in S1 Appendix.

In order to give some intuition, we consider the following special cases, where the SSD assumes a simpler form.

#### Special case 1

Suppose that the random vectors of the observation functions are independent and identically distributed, i.e., 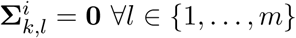 and *k ≠ l*. In this case, the *k*-th element of ***π***_*i*_ is

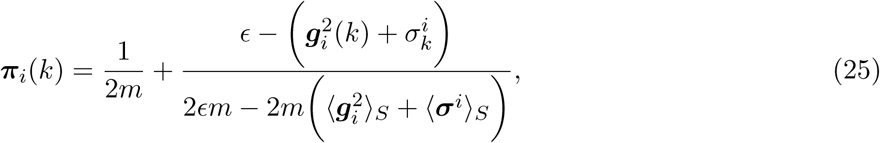

where

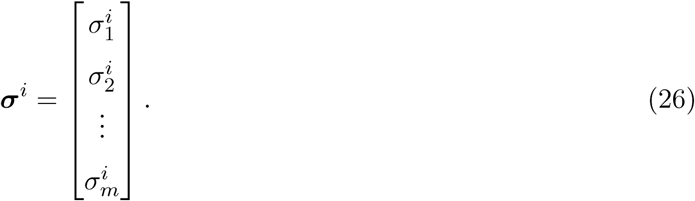

Note that a small value is assigned to ***π***_*i*_(*k*) if the weighted heterogeneity ***g***_*i*_(*k*) is large. In contrast, a large value is assigned to ***π***_*i*_(*k*) if ***g***_*i*_(*k*) is small. As a consequence, ***π***_*i*_(*k*) carries the heterogeneity information of the observations.

#### Special case 2

When the kernel scale *ϵ → ∞*, the information about the heterogeneity of each observation is lost, since the same weights are assigned to all the edges. As a consequence, ***π***_*i*_ becomes just a constant vector, given by

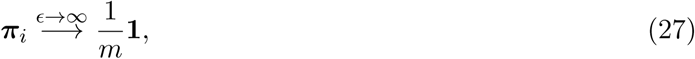

where **1** ∈ℝ^*m*^ is an all-ones vector.

### Binary Hypothesis Testing

Suppose that 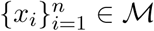 are realizations of a random variable *X*, which follows a bimodal distribution stemming from two hypotheses: ℋ_1_ and ℋ_2_; ℋ_1_ has probability *α* and ℋ_2_ has probability (1 − *α*), where 0 *< α <* 1. Denote the set of data points from hypothesis ℋ_1_ by Ω_1_ and the set of data points from hypothesis _2_ by Ω_2_. Recall that for each data point *x*_*i*_, *f*_*j*_(*x*_*i*_) are the realizations of the elements of the random vector *V* ^*i*^. Since *V* ^*i*^ depends on the random variable *X*, assume that *V* ^*i*^ also follows a bimodal distribution, which is induced by the bimodal distribution of *X*. Particularly, consider a Gaussian setting, where *V* ^*i*^ is sampled from *𝒩* (***m***_1_, **Σ**_1_) with probability *α* and from *𝒩* (***m***_2_, **Σ**_2_) with probability of (1 − *α*). Similarly, assume that the random variable 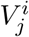 follows a bimodal distribution: sampled from 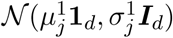 with probability *α* and from 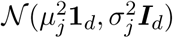 with probability (1 − *α*), where the respective probability density functions are 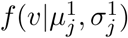 and 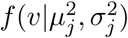.

A naïve approach for binary hypothesis testing would be to directly compare the densities of the two hypotheses for each observation separately. Particularly, based on the realizations from only one observation function *f*_*j*_, the average probability of error attained in a Bayesian setting with the MAP estimator [28] is given by

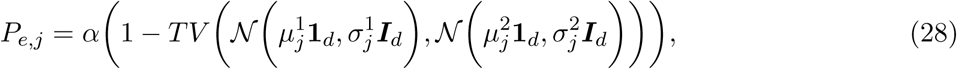

where 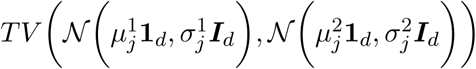 denotes the total variation between the realizations of 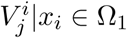 and 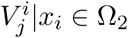, defined by

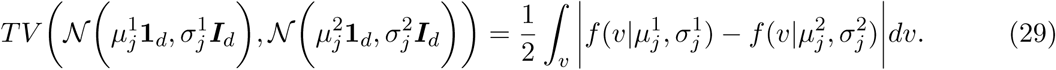

According to [11], consider 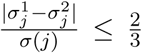, where 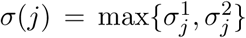. The total variation above between two Gaussian distributions is bounded by

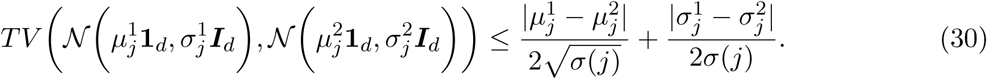

We seek another more discriminative approach for binary hypothesis testing. For this purpose, we propose a method based on the SSDs. Since the obtained SSDs represent probability distributions, the average probability of error is given by

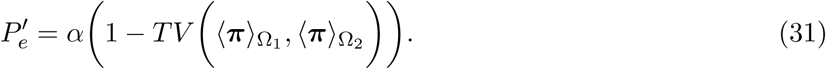

According to Proposition 1, the total variation between two SSDs associated with data points from two hypotheses can be explicitly expressed as the *l*_1_ distance given by

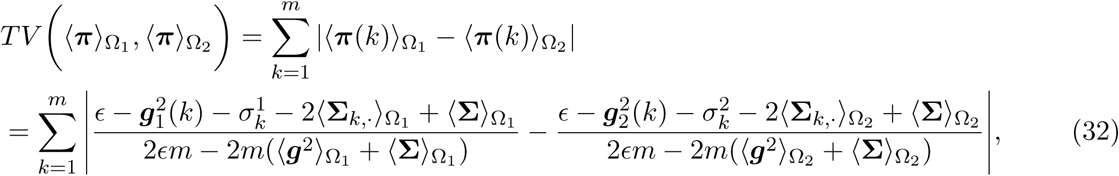

which consists of three main components: the variances 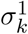 and 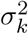, the weighted heterogeneities ***g***_1_ and ***g***_2_, and the covariance **Σ**.

The total variation of the measurements in Eq (29) and the total variation between the SSDs in Eq (32) can be used to distinguish between the two hypotheses. In the following, we specify the conditions, under which the total variation based on the SSDs in Eq (31) is larger, and hence, leading to smaller error compared to the standard MAP estimator using a single observation as specified in Eq (28).

#### Proposition 2

*Suppose* 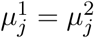 *and* 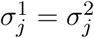, *which imply by definition (or by Eq (30)) that the total variation between the distributions corresponding to the two hypotheses is zero, i.e*.,

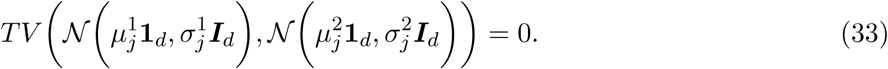

*This means that not only the standard MAP estimator but also any estimator based directly on single channel observations cannot distinguish between the two hypotheses. Conversely, from Eq (32), the SSDs may carry a distinction capability, that is*,

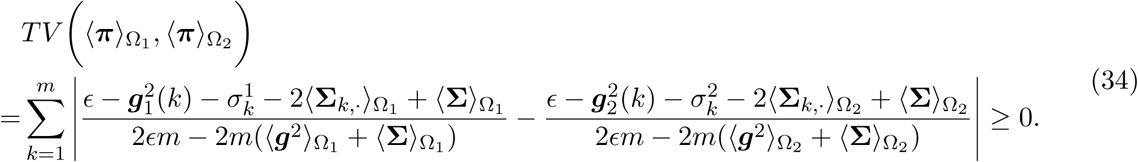

Proposition 2 demonstrates that there are cases where a single observation cannot be used for distinguishing between the two hypotheses. However, in such cases, the SSDs may enable us to distinguish the hypotheses due to possible differences in either the heterogeneity or the covariances. Proposition 2 is further demonstrated in the context of the localization toy problem in Simulation #2.

To further expand the analysis, we make the following assumptions.

#### Assumption 1

*The empirical mean of the weighted heterogeneity is approximately the same under the two hypotheses:*

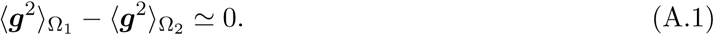

#### Assumption 2

*The empirical mean of the covariance matrices is approximately the same under the two hypotheses:*

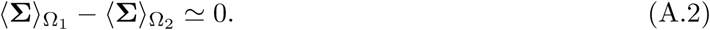

Note that if Assumptions (A.1) and (A.2) hold, implying that 2*ϵm* − 2*m*(⟨**g**^2^⟩_Ω1_ + ⟨**Σ**^2^⟩_Ω1_ ⋍ 2*ϵm* − 2*m*(⟨**g**^2^⟩_Ω2_ + ⟨**Σ**^2^⟩_Ω2_, then the *l*_1_ distance between the SSDs in Eq (32) can be recast as

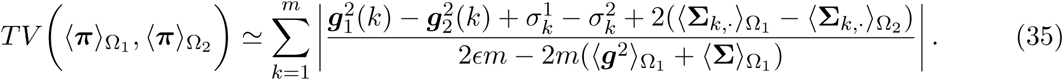

#### Proposition 3

*Suppose that Assumptions (A.1) and (A.2) hold. Suppose* 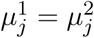, *which implies that the upper bound of the total variation at the j-th element in Eq (30) only depends on the variance*

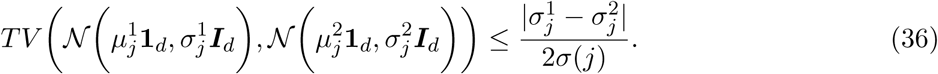

*In addition, suppose that (i) the covariance between the jth observation and the other observations under the two hypotheses is approximately equal* 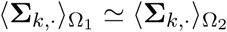, *(ii) the empirical mean of the difference of variance and weighted heterogeneity of the two hypotheses is greater than the difference of j-th variance, i.e*., 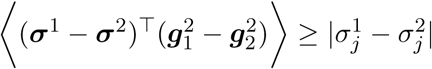, *and (iii) the weighted heterogeneity of ℋ*_1_ *is sufficiently large such that* 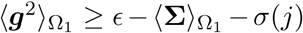. *Then, the l*_1_ *distance between the SSDs can be recast and bounded from below by*

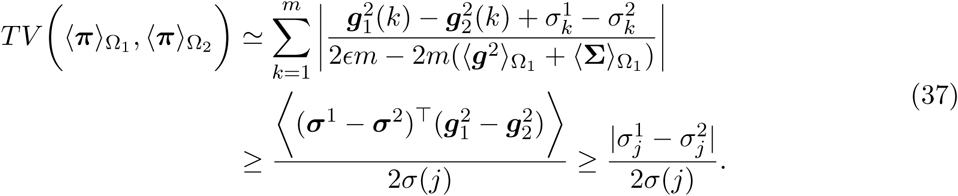

*It follows that*

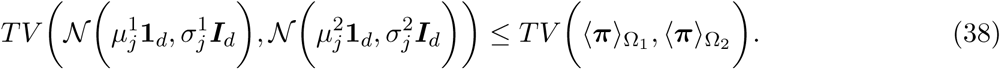

This proposition implies that when the assumptions hold, the probability of error based on SSD, which indirectly takes into account the mutual-relations between all observations, facilitates a better distinction of the two hypotheses compared to the standard MAP estimator computed from the best sensor. This property is further demonstrated in the localization toy problem in Simulation #3.

#### Proposition 4

*Suppose the conditions of Proposition 3 hold. In addition, suppose that the random vectors of the observations are independent and identically distributed, i.e*., 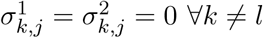 *then*

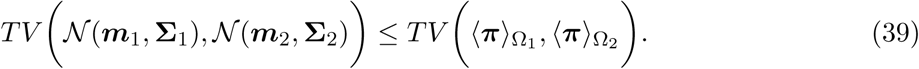

This proposition shows that the SSDs enable us a better distinction between the two hypotheses compared to the MAP estimator based on the distributions of the multi-feature observations. In other words, the heterogeneity comprising the SSD has a significant contribution to the ability to recover the information about the latent data points 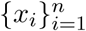 on the underlying manifold ℳ, thereby leading to accurate binary hypothesis testing.

### Imaging mass cytometry (IMC)

IMC is a relatively new imaging method, which enables to examine tumors and tissues at subcellular resolutions, giving rise to images consisting of the intensities of multiple proteins [5, 15, 7]. Our goal is to identify the sensitivity of lung cancer subjects to treatment with PD-1 axis blockers, given their IMC multiplexed observations. More specifically, we aim at a binary prediction task: identifying whether the subjects responded or did not respond to the treatment.

We analyze two IMC datasets consisting of baseline/treatment tumor samples from non-small cell lung cancer subjects profiled with 29 markers^1^ representing phenotype and functional properties of both tumor and immune cells. The study was approved by the Yale University Human Investigation Committee protocols #9505008219 and #1608018220; or local institutional protocols which approved the patient consent forms or, in some cases waiver of consent when retrospectively collected archive tissue was used in a de-identified manner.

The resolution of the IMC images is 1 *μm*^2^ per pixel. These subjects received treatment with PD-1 axis blockers. Based on the clinical sensitivity to the treatment, the subjects are categorized as: *responders* and *non-responders*. A standard pre-processing using z-score normalization is applied to each marker. We remark that the mean and the standard deviation are computed based only on pixels in which the marker has non-zero values.

Our analysis does not consider the entire image, but rather focuses on ROIs with the highest Cytokeratin expression levels, as Cytokeratin is expressed only in the tumor cells. In our experimental study, the ROIs are image patches of size 35×35 pixels, and we select 14 ROIs per sample by searching the patches with the maximal mean value of Cytokeratin. The size of the patch and the number of ROIs were determined empirically to yield maximal performance with cross-validation.

Using the present work notation, the intrinsic representation of the information embodied at each ROI is denoted by *x*_*i*_, which is assumed to be a data point from some hidden manifold ℳ. Our working hypothesis is that the distribution of 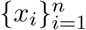 on the hidden manifold ℳ is bimodal, which is induced by the sensitivity of the subjects to the treatment; data points *x*_*i*_ at ROIs (patches) within tissues of responders are located in one region of the manifold, and data points *x*_*i*_ at ROIs (patches) within tissues from non-responders are located in another region of the manifold. Given a data point *x*_*i*_, our two hypotheses, ℋ_r_ and ℋ_n_, are whether *x*_*i*_ is a realization from the distribution of responders or non-responders, respectively. Next, recall that the intrinsic representation *x*_*i*_ is hidden. Instead, the observations are the expression levels of the markers at the ROIs (patches), which are represented mathematically by the observation functions *f*_*j*_(*x*_*i*_) ∈ℝ^*d*^ for *j* = 1, …, 29 whose domain is the hidden intrinsic manifold ℳ and range is an image patch of *d* = 35 × 35 pixels corresponding to the expression level of marker *j* at the ROIs associated with *x*_*i*_. Collecting the patches corresponding to all the markers at a specific ROI *x*_*i*_ gives rise to the set of multi-feature observations 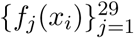. In Fig 2, we illustrate the IMC multiplexed observations from a single subject and the set up under consideration.

**Figure 2:**
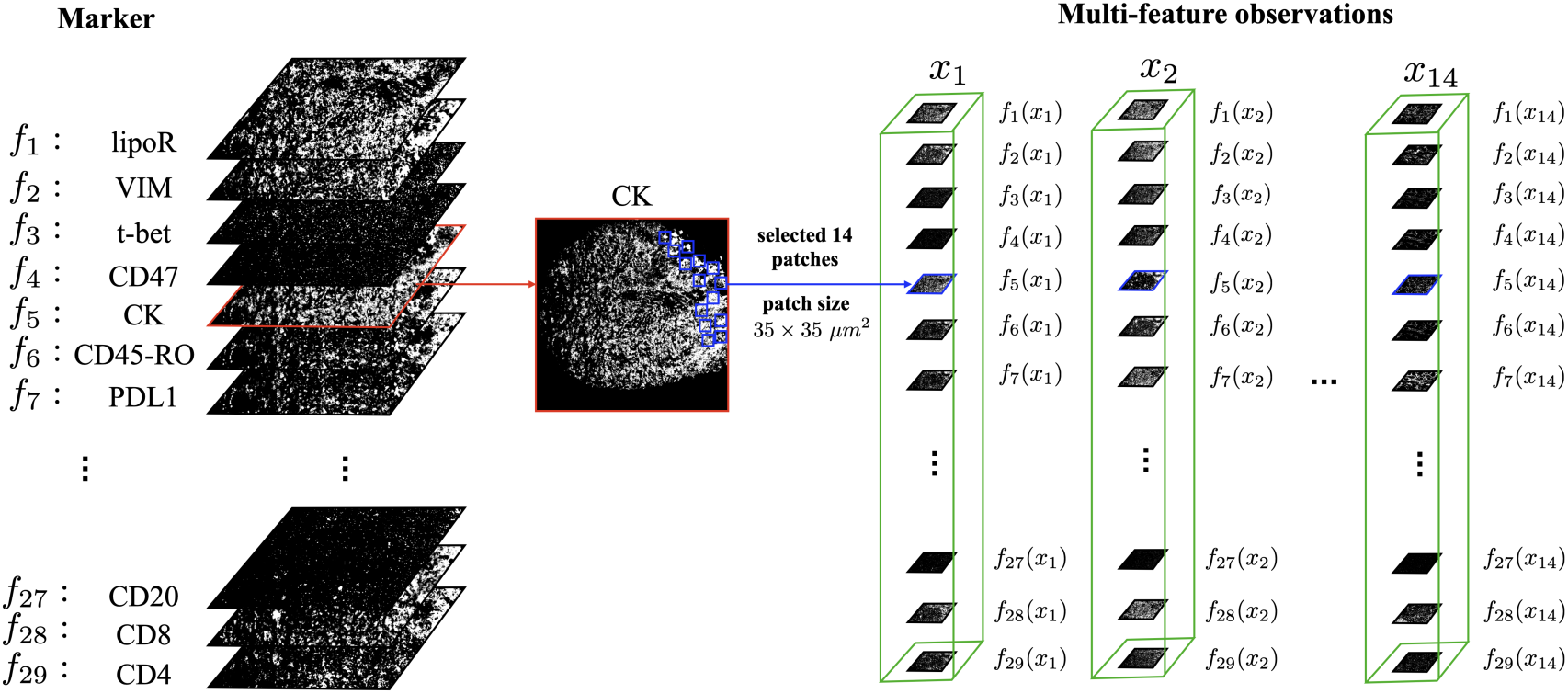
An illustration of the IMC multi-feature observations from a single subject. In the IMC data of a single subject, we focus first on the Cytokeratin marker, which is used as an indicator of tumor cells. We select 14 ROIs positioned at *l*_*i*_ for *i* = 1, …, 14. These ROIs are patches of size 35 × 35 *μm*^2^ with the highest Cytokeratin expression levels. We assume that these ROIs have intrinsic hidden states represented by *x*_*i*_. The expression of all 29 markers at these ROIs are viewed as the multi-feature observations 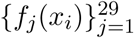 for *i* = 1, …, 14.

The identification of the subject’s response to treatment from the IMC data is based on the application of Algorithm 1. This algorithm fits well the problem at hand due to the following main reasons. First, we assume that the true intrinsic information about the tissues, enabling us to distinguish between the two hypotheses, is captured by the mutual-relationships between the markers, which are extracted by Algorithm 1. Second, directly comparing observations from the different markers, *f*_*j*_(*x*_*i*_), is inapplicable since each ROI comprises different cells and different tissue structures. Algorithm 1 circumvents this problem by computing an intrinsic signature of the observations. Third, due to the different dynamic range of the nominal values of the observations from the different markers, a naïve concatenation of the multi-feature observations is inadequate (in contrast to the localization example).

We apply Algorithm 1 to the sets of observations, 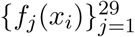 for *i* = 1, …, *n*. Then, we apply to its low-dimensional output an RBF SVM classifier with a leave-one-subject-out (LOSO) cross-validation. We note that the hyper-parameters of the algorithm, namely, the patch size, the number of patches per subject, and the scale of the kernel, were chosen so that optimal empirical results are obtained. Importantly, we report that the performance of the LOSO cross-validation was not sensitive to the particular choice of parameter values. In order to assess the classification performance for each subject, we compute the average of the classification results of all the ROIs of that subject. We compare the SSD-based representation by Algorithm 1 to other representations obtained by two competing algorithms. The first is a direct application of diffusion maps to the sets of multi-feature observations 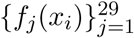 for *i* = 1, …, *n*. The second is HKS [35]. The dimensions of representation obtained by Algorithm 1 and by the two competing algorithms are determined by a variant of the Jackstraw method, described in Section Determining the dimension of data. In order to visualize the representations obtained by the different methods, we use t-SNE [25].

#### Dataset 1

Dataset 1 consists of 26 responders and 29 non-responders, so that the chance level of classifying a patient as responder is 47.27%. Overall, we extract from this dataset a total of (26 + 29) × 14 = 770 ROIs (patches).

Considering all 29 markers, the low-dimensional representation obtained by Algorithm 1 consists of 7 dimensions, leading to a total LOSO classification error of 9.1%, and the corresponding AUC is 0.909. In the direct application of diffusion maps to the multi-feature observations, the number of dimensions is set to 13 and the LOSO classification error is 41*/*8%, and the AUC is 0.582. By using HKS as a feature vector, the number of dimensions is set to 7, resulting in LOSO classification error of 25.5% and the AUC is 0.745.

Fig 3 presents the 3D t-SNE visualization of the representations colored by the response status and by the subjects index. Fig 4 presents the confusion matrix of the LOSO classifications. Evidently, the classification obtained by Algorithm 1 is superior compared to the other two competing methods. In addition, in the t-SNE visualization, we observe that the unsupervised separation of the patches from responders and non-responders is most pronounced in the low-dimensional representation obtained by Algorithm 1.

**Figure 3:**
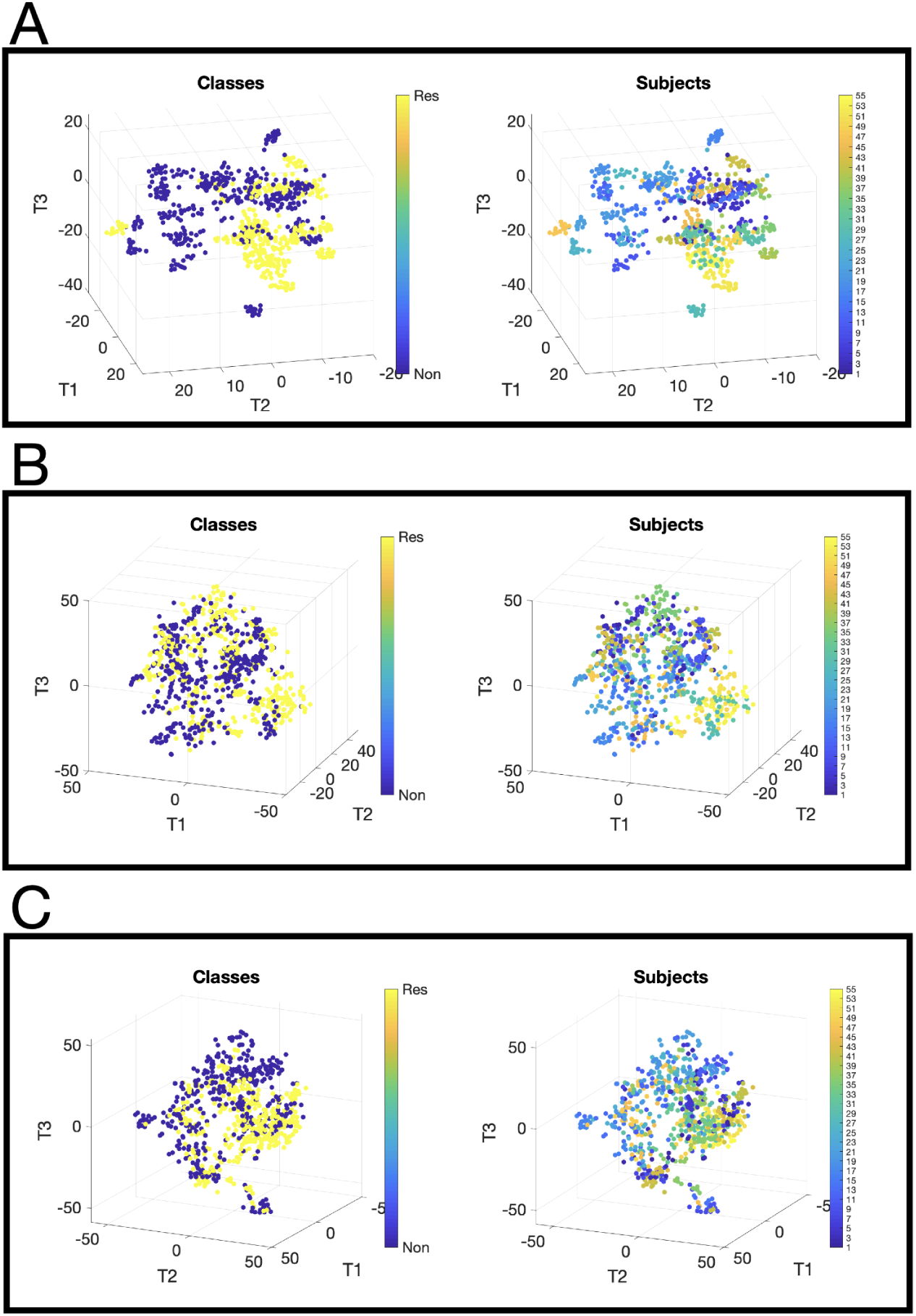
t-SNE plots of ROIs response status and patient ID for Algorithm 1, DM and HKS. The columns, from left to right, show the visualization of the patches by t-SNE colored by response status and the visualization of the patches by t-SNE colored by the index of the subjects. The rows, from top to bottom, depict the obtained visualizations of Dataset 1 by Algorithm 1, diffusion maps applied directly to the multi-feature observations, and using HKS as the feature vector.

**Figure 4:**
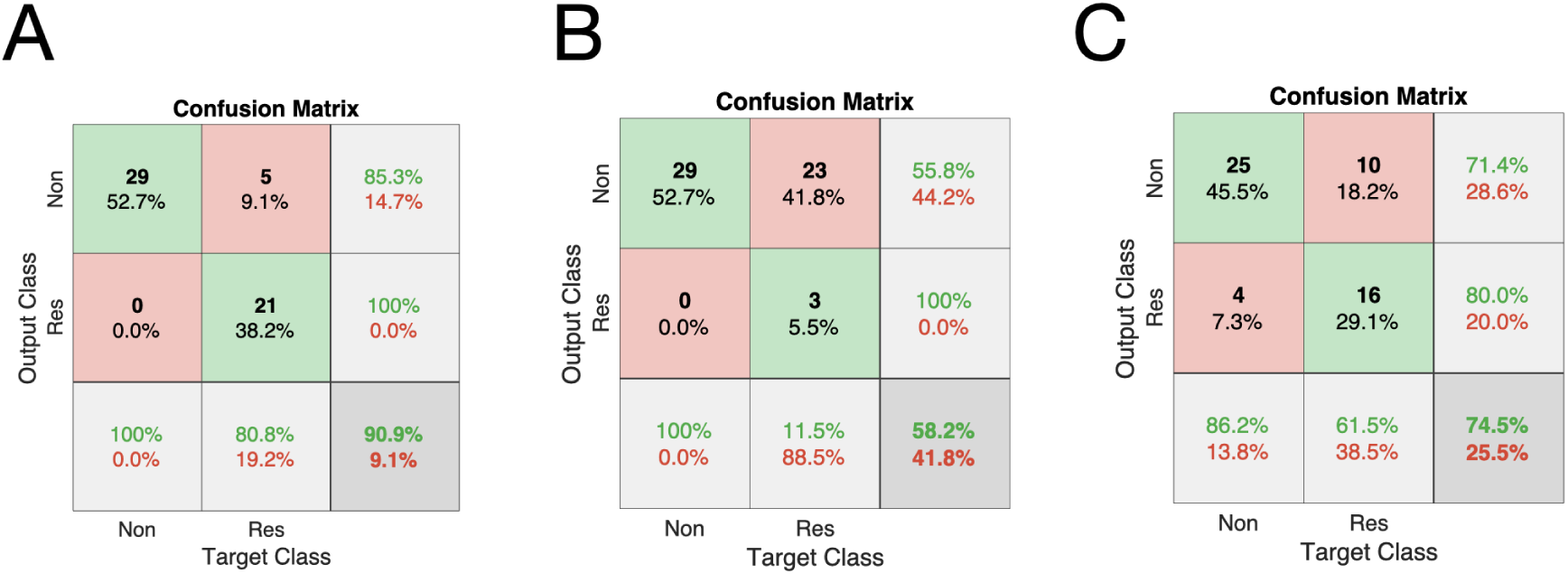
Confusion matrices of Dataset 1 obtained by Algorithm 1, DM and HKS. The confusion matrix of the classification to response status obtained by a leave-one-subject-out cross-validation using an RBF SVM classifier. (A) by Algorithm 1, (B) diffusion maps applied directly to the multi-feature observations and (C) using HKS as the feature vector.

#### Dataset 2

Dataset 2 consists of 11 responders and 18 non-responders, thus, the chance level of classifying a patient as responder is 37.93%. A 3 × 3 median filter is applied to each marker image as a pre-processing for denoising. Compared to Dataset 1, the images of Dataset 2 are denser and allow for the application of the median filter. Overall this dataset has a total of (11 +18) × 14 = 406 ROIs (patches).

The low-dimensional representation obtained by Algorithm 1 consists of 10 dimensions retained by the variant of Jackstraw procedure, leading to LOSO classification error of 6.9%, and the AUC is 0.931. In the direct application of diffusion maps to the multi-feature observations, the number of dimensions is set to 28, yielding LOSO classification error of 17.2%, and the AUC is 0.828. The dimension of diffusion maps with the HKSs is set to 7, resulting in LASO classification error of 17.2%, and the AUC is 0.828.

Similar to Fig 3 and Fig 4, Fig 5 shows the visualizations obtained for Dataset 2 and Fig 6 presents the corresponding confusion matrix. The results indicate that the classification obtained by Algorithm 1 is superior compared to the two competing methods. In addition, the t-SNE visualizations show that in the representation obtained by Algorithm 1, in a completely unsupervised manner, the separation of the ROIs (patches) into two clusters according to the response status is evident.

**Figure 5:**
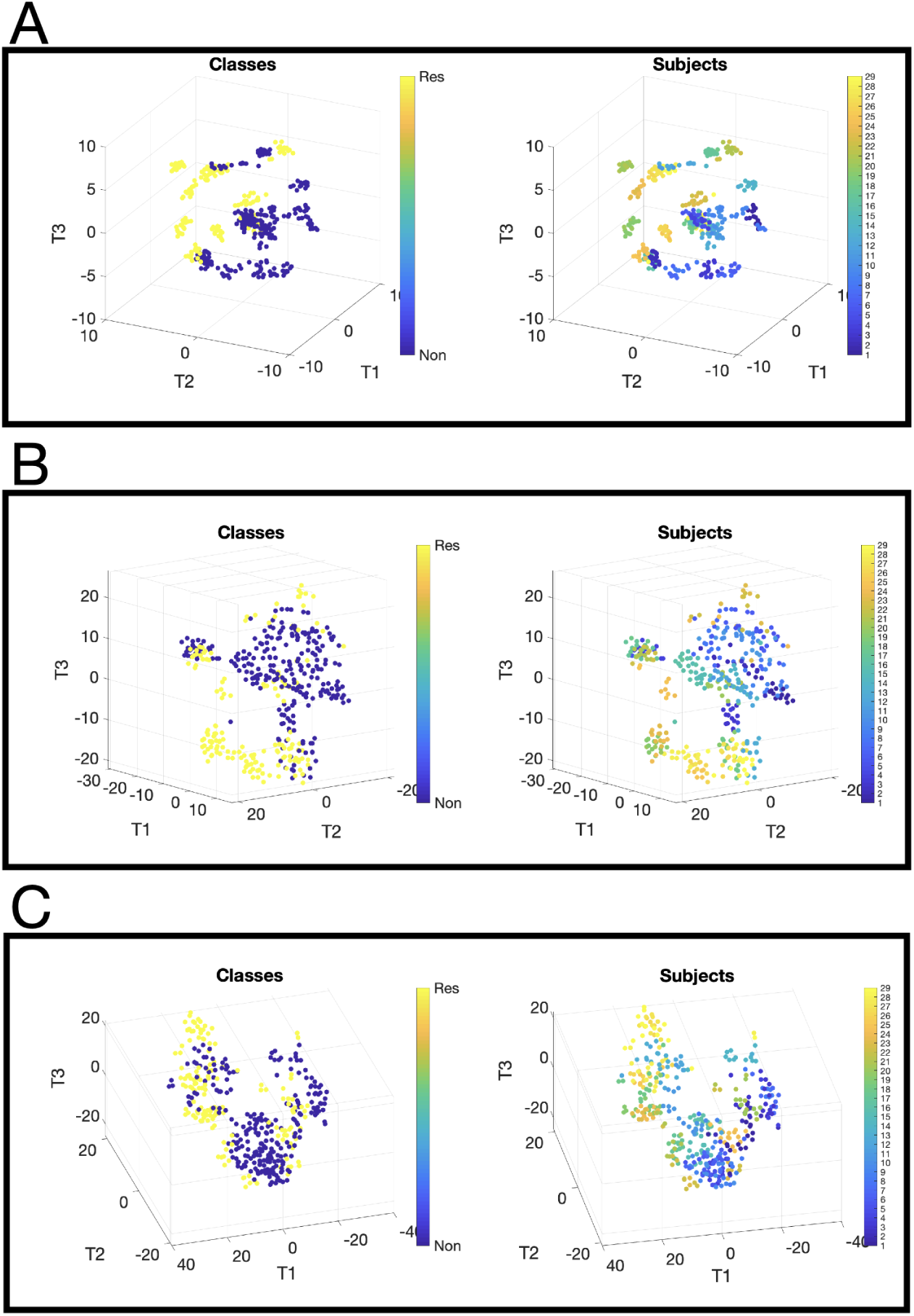
t-SNE plots of Dataset 2. Same as Fig 3 but for Dataset 2.

**Figure 6:**
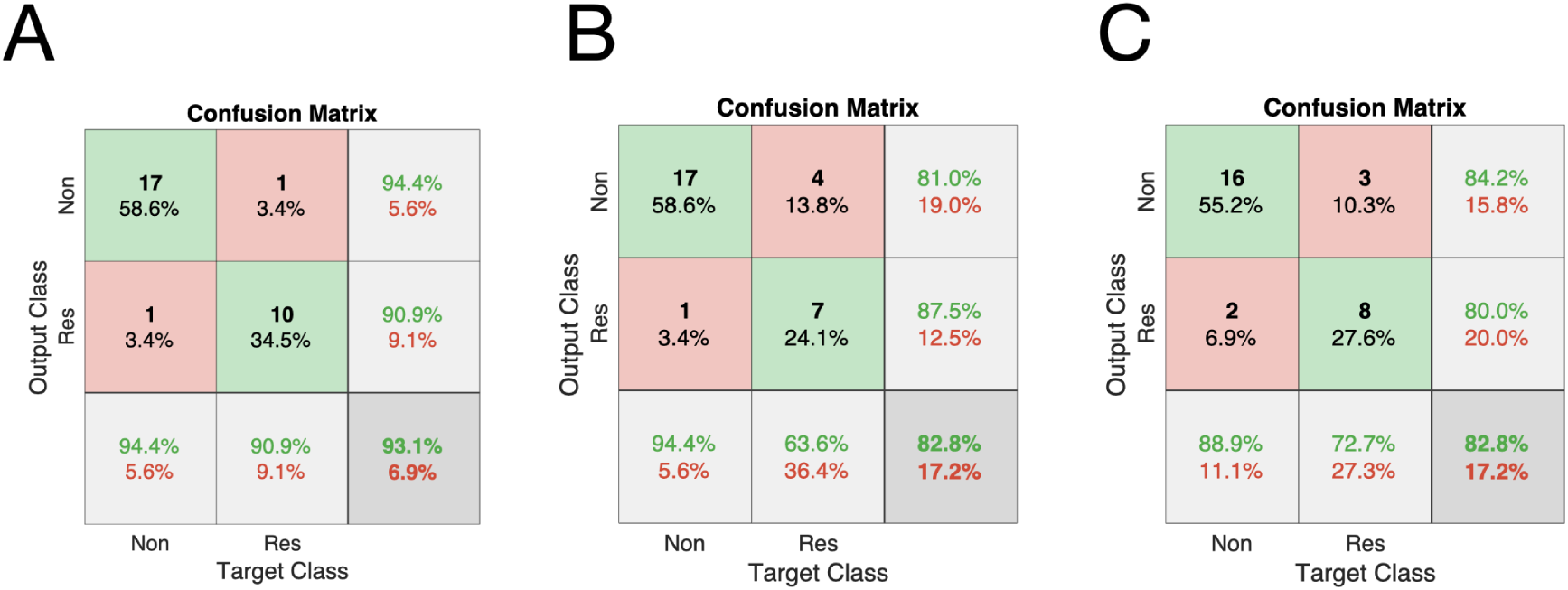
Confusion matrix of Dataset 2. Same as Fig 4 but for Dataset 2.

## Discussion

We presented a two-step graph analysis approach. The first step is applied to the multi-feature observations of the data points, where their mutual-relationships are extracted. This step is implemented by computing the SSD of a random walk defined on the graph whose nodes are the observations. The resulting SSD can be viewed as a signature or a characteristic vector of the data point and is analogous to traditional signatures from other domains, such as the heat kernel signature (HKS) [35] in the field of shape analysis and PageRank [29] in webpage ranking. The second step is applied to the signatures obtained at the first step, for the purpose of constructing an intrinsic low-dimensional representation of all the data points.

Previous attempts to analyze such mulitplexed datasets involve various approaches, including direct comparisons of the marker expressions, the cell morphology, and interactions in cell neighborhoods, to name but a few [2]. Our method introduces a new approach, building a new representation of the multiplexed data in two steps. Since each of the steps involves a construction of a graph, the entire procedure can be viewed as building a graph of graphs.

While the algorithm is described in a general setting of multiplexed data fusion, our theoretical analysis is focused on binary hypothesis testing. In comparison with a traditional statistical estimation approach, we show that our method exhibits advantages, implying that the mutual-relationships between the multi-feature observations are well captured. In the context of IMC, this could minimize the effect of deviation in individual marker scores and cell/tissue heterogenenity.

We apply the proposed method to IMC data and show that solely from the imaging data, we can distinguish between two different sensitivity levels to treatment. Since our approach does not rely on rigid prior knowledge or access to labels, it has the potential of identifying biological relevance of novel parameters or marker patterns in treatment responses by analyzing dominant factors contributing more to the model stratification. Importantly, we remark that in contrast to common practice, the proposed approach does not require cell-segmentation as a precursor.

It is conceivable that the most important hyperparameter of our method is the scale parameter. In S1 Fig, we present a toy example demonstrating that different values of the scale parameter *ϵ* lead to multiscale signatures capturing local and global features. We demonstrate that different scales facilitate the extraction of different features of the data. In future work, we plan to further explore the role of the scale and to devise multiscale signatures. Another possible direction for future research relies on the fact that our method is general and can be extended to other multiplexed datasets, for example, Slide-seq [31], High-Density Spatial Transcriptomics [40, 39], MIBI-TOF [17], and DBiT-seq [22].

## Materials and methods

### Diffusion maps

Manifold learning is a class of nonlinear techniques that embeds high dimensional data points into a low dimensional space, relying on the assumption that the high-dimensional data lie on a low dimensional manifold ℳ [3, 10, 32]. In order to “learn” the manifold from a discrete set of data points, a graph is typically defined, where the graph nodes are the data points and the edges are determined according to some similarity notion. Since the manifold information is entirely captured by its Laplacian, the discrete counterpart, the graph Laplacian is used to build a low-dimensional embedding that respects the manifold in some sense [4]. To this end, common practice is to compute and exploit the spectral decomposition of the graph Laplacian. Diffusion maps is one of these methods, which constructs a random walk on the graph and represents the data points in a low-dimensional space preserving the neighborhood information [10].

Consider a set of data points 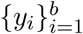, where *y*_*i*_ ∈ℝ^*d*^ for *i* = 1, *…, b*. An undirected weighted graph *𝒢* = (*𝒱, ε*, ***W***) is constructed from the data points, where the vertex set is *𝒱* = (*y*_1_, *y*_2_, *…, y*_*b*_) and the weights of the edges connecting two vertices are determined by a measure of similarity between any two data points, e.g., by

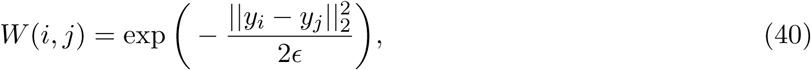

where *i, j* ∈ {1, *…, b}* and *ϵ >* 0 is a scale parameter. Common practice is to set *ϵ* as the median of the distances between the graph nodes. Note that *ϵ* implicitly induces a notion of locality: it can be viewed as the (squared) radius of the neighborhood around each node, so that only nodes within this radius are considered as neighbors in the graph.

Next, a random walk ***P*** on the data points is constructed by normalizing the weight matrix ***W***

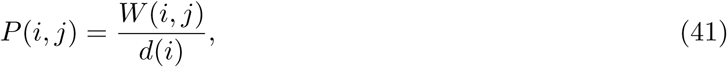

where 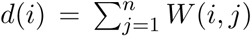. ***P*** is the transition matrix of a Markov chain defined on the data points 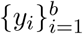 (graph vertexes), where the entry *p*(*i, j*) describes the probability of a random walk transitioning from the node *y*_*i*_ to the node *y*_*j*_ in a single step. Raising the transition matrix ***P*** to a power *t* can be viewed as applying the Markov chain to the data points *t* times.

Since ***P*** is similar to a symmetric and positive-define matrix, ***P*** has a biorthogonal right- and left-eigenvectors 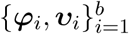 with the eigenvalues 1 = *λ*_1_ *≥ λ*_2_ *≥* … *≥ λ*_*b*_ = 0. Consequently, the spectral decomposition of ***P*** ^*t*^ is given by

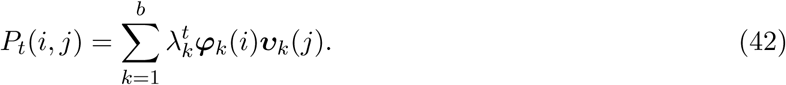

The diffusion distance 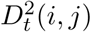 between two data points *y*_*i*_ and *y*_*j*_ in the data set is defined by

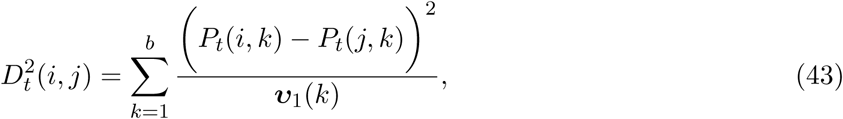

which measures the similarity of two points based on the evolution of their probability distributions, and depends on all possible paths of length *t* in the graph between any two points. Namely, if two points are connected by a large number of paths, then the diffusion distance between them will be small. Conversely, if there are only few paths connecting two points, then the diffusion distance between them will be large.

The diffusion maps is defined by [10]

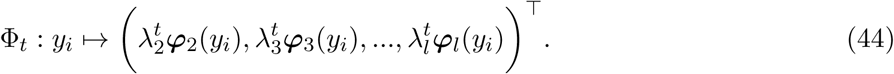

We remark that in many cases, due to the typical fast decay of the eigenvalues of ***P*** ^*t*^, *l* can be set to be smaller than *d*, thereby achieving dimension reduction. In addition, ***φ***_1_ is a constant vector and therefore is not used in the mapping.

It can be shown that the diffusion distance can be approximated by the eigenvalues and eigen-vectors by [10]

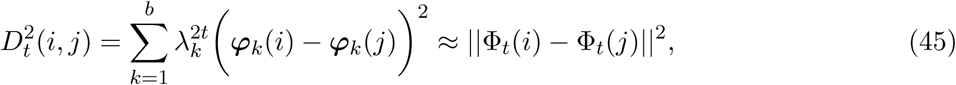

where equality is reached for *l* = *b*. Namely, the diffusion distance can be approximated by the Euclidean distance between the diffusion maps of the data points.

### Determining the dimension of data

A common problem in diffusion maps setting is how to choose the dimension *l*. The authors in [9] proposed *Jackstraw* to identify the number of principal components (PCs) in the context of principal component analysis (PCA) [13]. We present here a variant of *Jackstraw*, adapting it to diffusion maps.

Given a random walk ***P*** constructed from a set of *b* data points, the associated eigenvalues are 1 = *λ*_1_ *≥ λ*_2_ *≥* … *≥ λ*_*b*_ with corresponding right-eigenvectors 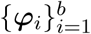. Collect the eigenvalues into a vector, denoted by ***λ***, where ***λ*** = (*λ*_1_, *λ*_2_, *…, λ*_*b*_)^*T*^. Let 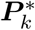 consist of the random permutation of ***P***. Apply eigenvalue decomposition to 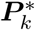 and obtain the corresponding eigenvalues vector 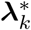. Repeat this shuffling procedure *s* times and obtain a set of vectors 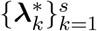. The dimension of the representations is determined by

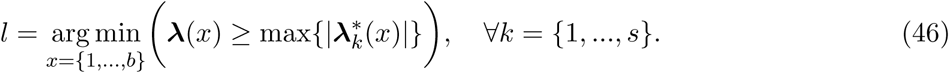

Note that the absolute values of the eigenvalues of 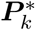 are considered because 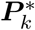 is not necessarily symmetric and therefore its eigenvalues 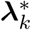 are not guaranteed to be real.

### Related work

#### Heat kernel signature

There are several shape analysis signatures obtained by spectral methods with different geometric properties such as isometry and deformation invariance [1, 30, 33, 35], related to the proposed method. One of the notable shape signatures is based on the heat diffusion on a shape, called Heat Kernel Signature (HKS) [35]. Broadly, the HKS is obtained by the eigenvalue decomposition of the heat kernel defined on the shape. In the context of our problem, since the heat kernel and the Laplace-Beltrami operator Δ_*i*_ share the same eigenbasis, and since the discrete graph Laplacian converges (point-wise) to the Laplace-Beltrami [26]

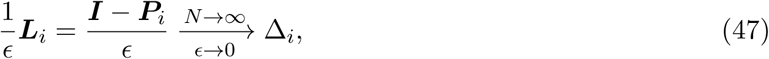

then, a discrete counterpart of HKS is given by

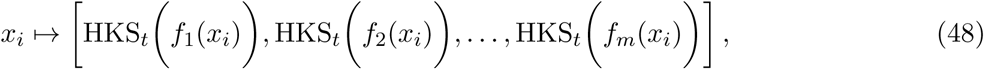

where

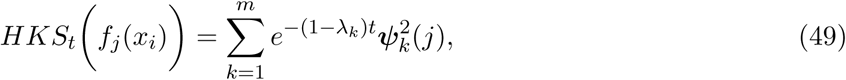

where *λ*_*k*_ and ***ψ***_*k*_ are the *k*-th eigenvalue and *k*-th eigenvector of the random walk, respectively. For more details on HKS, we refer the readers to [35].

Similarly to the DKS in Eq (12), the HKS can also be viewed as a low-pass filter. Observe that, for small *t*, the HKS approximates the DKS by Taylor expansion; for other *t* values, the weights assigned by the HKS decay faster than the weights of DKS, and therefore, the DKS gives more attention to finer structures.

#### Nodes ranking

In the context of web page ranking, there are several traditional algorithms based on the spectral analysis of directed graphs. Among them, the celebrated PageRank score is based on the stationary distribution of a random walk representing the popularity of linked web pages [29]. Hyper induced topic search (HITS) is another related algorithm, identifying the influential nodes using a random walk on a graph. There, the graph nodes are the web pages, which are divided into two groups: authorities and hubs [18]. Both algorithms address the problem of web page ranking, where PageRank depends on the incoming links whereas HITS focuses on the outgoing links.

### Localization toy problem

To illustrate the challenge in the problem setting and the generality of the proposed solution, we present three simulations of different localization problems.

#### Simulation #1

Consider 800 objects on a 2-sphere in ℝ^3^ that can be located at four different regions. Each region consists of 200 objects. The positions of the objects are measured by 5 sensors, giving rise to the following set of observations 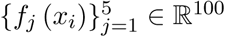, where *j* is the index of the sensor and *i* is the index of the object (position). Each sensor measures the position in *d* = 100 coordinates in the following way

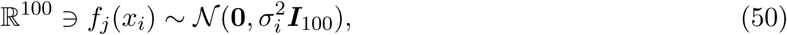

where the standard deviation of the measurement depends on the distance between the position of the sensor and the position of the object 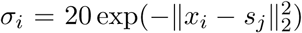 is the identity matrix of size 100 × 100, ∥ ·∥_2_ denotes the Euclidean norm and *s*_*j*_ denotes the 3D position of sensor *j* ∈ {1, …, 5} In other words, each object position is captured by *d* = 100 realizations of a Gaussian random variable with variance that is proportional to the distance between the sensor position and the object position. Note that the positions are captured by the sensor through the variance, therefore, they are difficult to infer directly by the multi-feature observations.

Fig 7 panel A, on the left-hand side, we illustrate the setting where the objects positions are depicted as dots, the different regions are marked by different colors (red, black, blue and yellow), and the sensors positions are marked by green stars. The right-hand side of Fig 7 panel A shows the (high-dimensional) sensor observations. Visually, it is evident that distinguishing between the different regions merely based on these observations is non trivial.

**Figure 7:**
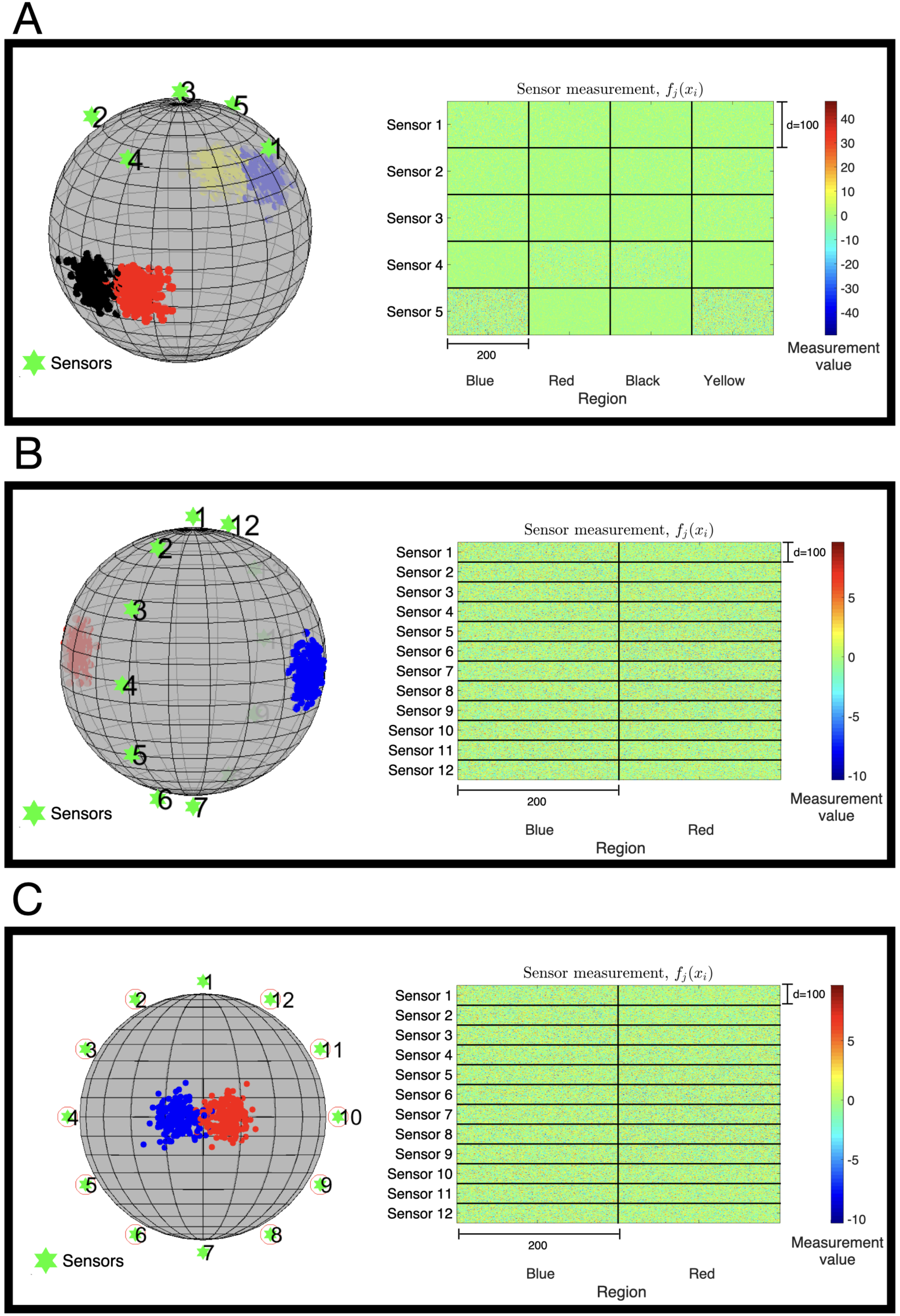
Illustration of localization toy problems. The first column depicts the objects location residing in different regions and the sensors locations. The second column demonstrates the multi-sensor observations. The rows, from top to bottom, show three different localization toy problems.

For illustration purposes, we view the problem as a classification problem, where given the high-dimensional multi-feature observations, the task is to identify in which region the object at *x*_*i*_ resides.

The observations from each sensor consisting of 800 object positions from four regions were processed in two stages: first, a 3D embedding is constructed by applying diffusion maps to the high-dimensional observations, and second, an RBF SVM is applied to the obtained embedding in order to classify the region. To evaluate the classifiers, we perform a 10-fold cross-validation. A similar two-step procedure is applied to the concatenation of the observations from all the sensors.

The performances obtained by the 10-fold cross-validations are presented. The percentages indicate the number of correctly classified positions divided by the true number of total positions in each class.

Table 1 presents the resulting classification accuracy, i.e., the number of correctly classified positions divided by the total number of positions in each region. The presented results are obtained using a 10-fold cross-validation. We observe that none of the sensors enables an accurate classification. Fig 8 sheds light on these poor classification results. The figure presents the diffusion maps 3D embedding of the multi-feature observations. The 3D embedded points are colored according to the four regions. We observe no particular clustering of the points by their respective region. Moreover, we show in Table 1 that a naïve concatenation of the observations from all the sensors do not yield a good classification either, which is visualized in Fig 8 panel A as well.

**Table 1:** Localization accuracy from measurements of Simulation #1.

| Region | Concatenated sensors | Sensor 1 | Sensor 2 | Sensor 3 | Sensor 4 | Sensor 5 |
| --- | --- | --- | --- | --- | --- | --- |
| Blue | 43% | 46% | 18% | 44.5% | 58% | 40.5% |
| Red | 54% | 15.5% | 52.5% | 45.5% | 48.5% | 55% |
| Black | 46.5% | 49.5% | 29% | 40% | 37.5% | 42% |
| Yellow | 38.5% | 33.5% | 52% | 30.5% | 42.5% | 50.5% |

**Figure 8:**
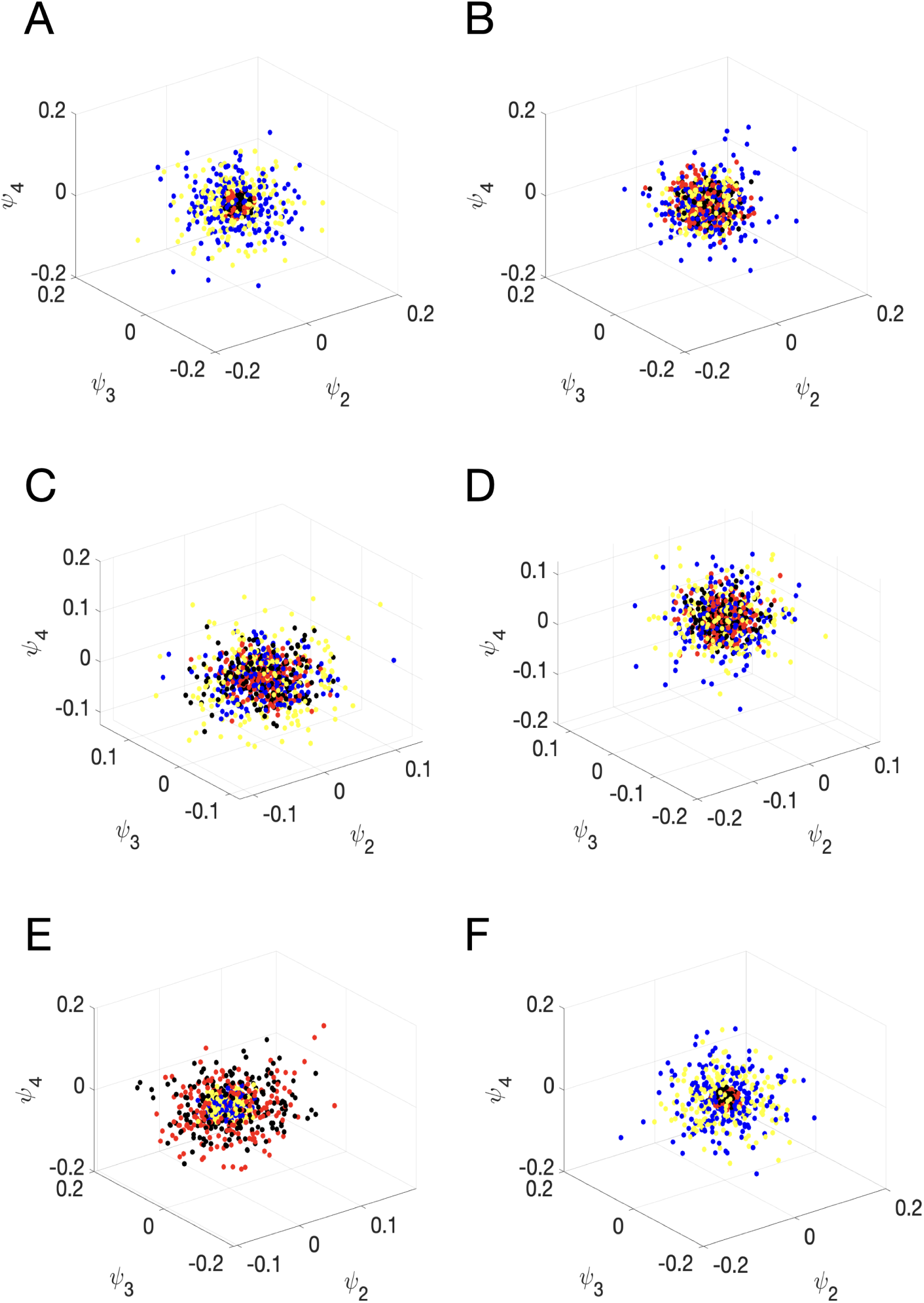
Diffusion maps directly applied to the observations in localization toy problems 1. The depicted embedded points are colored according to the region. A: The concatenated observations. B: The observations of Sensor 1. C: The observations of Sensor 2. D: The observations of Sensor 3. E: The observations of Sensor 4. F: The observations of Sensor 5.

Seemingly, in order to mitigate the problem, we could simply represent the objects positions by a vector of the variance of the observations. However, it would require prior knowledge about the sensing model, whereas our approach is model-free.

Fig 9 shows the results of the application of Algorithm 1 to the observations.

**Figure 9:**
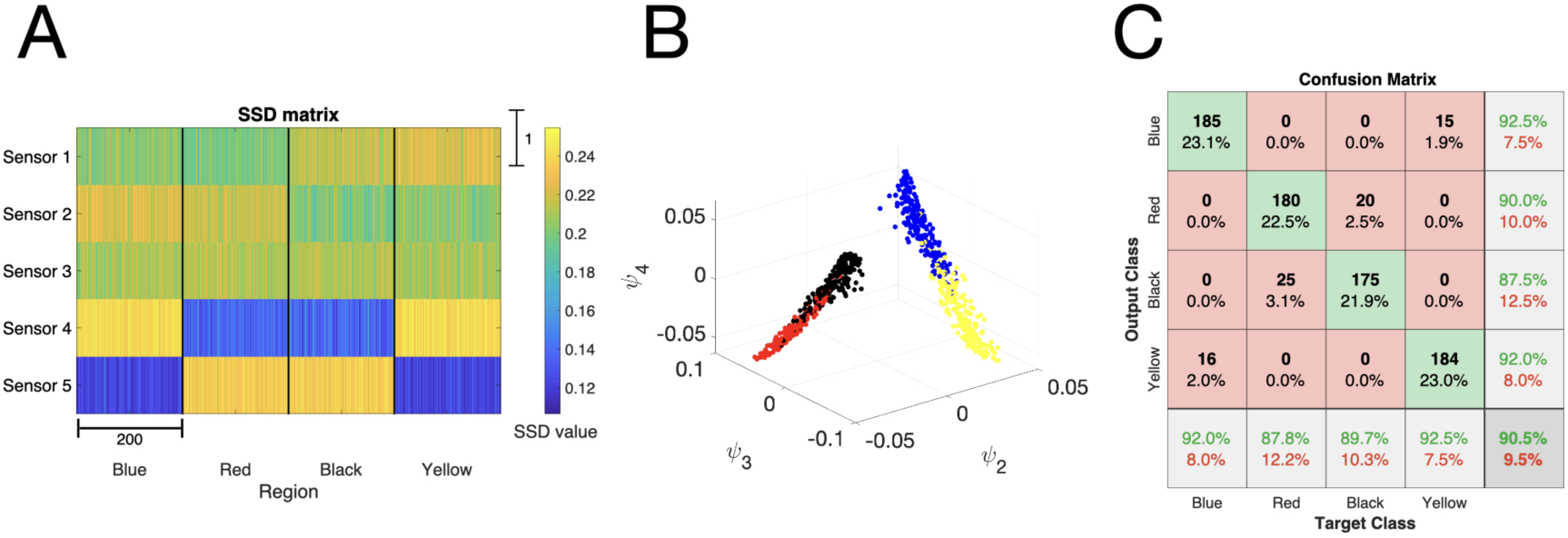
Application of Algorithm 1 to Simulation #1. A: The corresponding SSDs obtained by the proposed method. B: Diffusion maps embedding of the proposed method. C: 10-fold cross-validation obtained by an RBF SVM classifier.

Fig 9 panel A presents the SSDs obtained from all the objects. In contrast to Fig 8, Fig 9 panel A highlights the different region locations with SSD signature and it is only mildly affected by the difference in the dynamic range visible in Fig 7 panel A.

Fig 9 panel B presents the diffusion maps embedding constructed from the SSDs, where we observe a clear separation between the four regions. Fig 9 panel C shows the confusion matrix of the classification obtained based on the representation in Fig 9 panel B. We observe that the proposed method leads to significantly better classification results compared to the results in Table 1.

Comparing the classification results in Fig 9 panel C with the classification results in Table 1, show the importance of taking into account the mutual-relationships between the sensor observations, rather than processing the nominal values of the observations directly, which in this case, give rise to correct identification of the four regions.

#### Simulation #2

The main purpose of this simulation is to demonstrate Proposition 2 from Binary Hypothesis Testing Section.

Consider *n* = 400 positions, 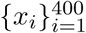, such that *x*_*i*_ ∈ *S*^2^ ⊂ℝ^3^, which are sampled from two different regions on the sphere. Suppose that the positions of the objects follow a bimodal distribution as depicted in Fig 7 panel B: an object is located in the blue region with probability 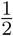 and in the red region with probability 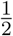. The two regions represent the two hypotheses, ℋ_1_ and ℋ_2_, where each region consists of 200 positions.

Here, we have 12 sensor observations 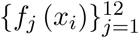, measuring the positions of the objects, located at *s*_*j*_. The multi-feature observations are random samples from

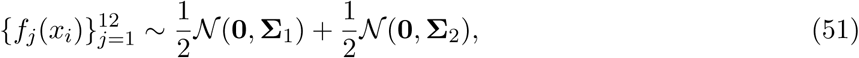

where *f*_*j*_(*x*_*i*_) ∈ℝ^100^, the standard deviation is 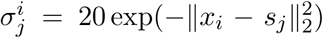, and the covariance Σ_*k,l*_ *∼* |*𝒩* (0, 0.5)| for 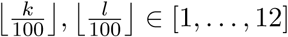.

In Fig 7 panel B, the object positions are depicted as dots, where the two regions are colored in blue and red, and the sensor positions are marked by green stars. Fig 7 panel B shows the high-dimensional sensor observations. The figure is divided to *m* = 12 vertical blocks and two horizontal blocks. Each block consists of 100 × 200 scalar observations corresponding to the observations of a single sensor from either the blue or red region, where *d* = 100 is the dimension of each observation and 200 is the number of the positions per region.

Note the symmetry in this setting, that is, the distances between the two regions and the sensors are approximately the same.

In this simulation, a direct computation can show that Assumptions *(A.1)* and *(A.2)* hold. In addition, the conditions of Special Case 1 are satisfied, and thus, the total variation of these sensor observations is zero. In other words, using a single sensor is insufficient to distinguish between the two regions. Conversely, we show that since the SSDs take into account the covariance information between the sensors, they allow us to make this distinction.

Fig 10 panel A shows a comparison between the classification results obtained by Algorithm 1 using SSDs and the classification results obtained by using the output of each sensor as well as the concatenation of the output from all the sensors. The performances are evaluated with a 10-fold cross-validation. We observe that the classification obtained by Algorithm 1 is significantly better than the classification obtained using the “raw” sensor outputs.

**Figure 10:**
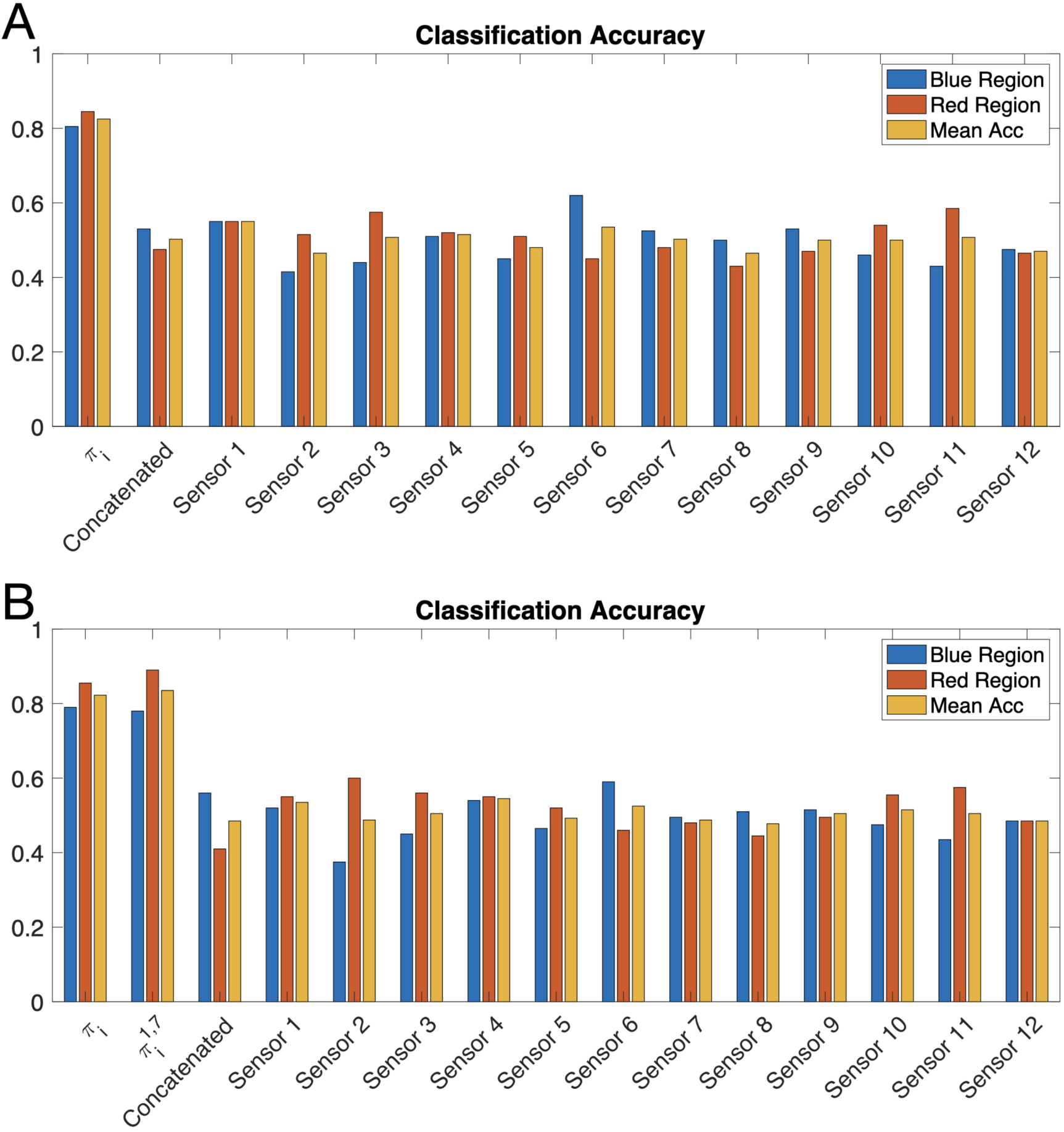
Localization accuracy obtained by Algorithm 1 for Simulation #2 and Simulation #3. The percentages indicate the number of correctly identified positions divided by the true number of total positions in each region. (A) Simulation #2 and (B) Simulation #3.

#### Simulation #3

This simulation is to demonstrate Proposition 3 from Binary Hypothesis Testing Section. Consider *n* = 400 positions, 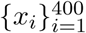, such that *x*_*i*_ ∈ *S*^2^ ⊂ℝ^3^, which are sampled from two different regions on the sphere, as depicted in Fig 7 panel C. The rest of the setting remains as in Simulation #2.

Note that here, only two sensor observations satisfy the conditions of Special Case 1, namely, 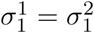 and 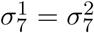, and thus, the total variation of only these two sensor observations is zeros.

For each data point, we compute the difference between the left-hand side and the right-hand side of the inequality in Proposition 2 per sensor. Then, in Fig 7 panel C, we circle the sensors attaining the largest difference. As expected, we observe that the circled sensors are positioned in non-symmetric orientations with respect to the locations of the two regions.

Similarly to the previous simulation, we compare the classification results obtained by Algorithm 1 to the results attained by applying the classification directly to the observations from each sensor separately and to the concatenation of the observations from all the sensors. In addition, we also compute the results of Algorithm 1 applied to all the sensors except Sensor 1 and Sensor 7, which are the least contributing according to the inequality in Proposition 2.

Similarly to Fig 10 panel A, Fig 10 panel B demonstrates the classification abilities of SSD. Specifically, we observe that by removing the least contributing sensors, namely Sensor 1 and Sensor 7, the recomputed SSDs 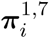 lead to slightly improved classification accuracy.

## Supporting information

## S1 Appendix. Detailed derivation of Proposition 1

### S1 Fig. Multi-SSD

We illustrate this multiscale property using a 3D shape from Princeton ModelNet40 database [41]. Suppose the points on the shape are the graph nodes, and compute the SSD of the graph with different values of *ϵ*, where the *ϵ* are chosen in logarithmic spacing between [10^−3^, 10^1.5^]. The color represents the values of SSD with different scales *ϵ* computed based on data points from a 3D shape of flowers in a vase. The red color represents high values of SSD, and, the blue color represents the low value of SSD. We observe that ***π***_*i*_ highlights junctions or hubs (in red), as in [24], both at local and global scales, depending on *ϵ*. We also observe that when *ϵ* is small, the SSD highlights the neck of each flower. When gradually increasing the value of *ϵ*, we observe that the SSD transitions toward the center of the shape, representing the global hub of the shape.

## Author Contributions

**Conceptualization:** Ya-Wei Eileen Lin, Tal Shnitzer, Ronen Talmon, Kurt Schalper, Yuval Kluger.

**Data curation:** Franz Villarroel-Espindola, Shruti Desai.

**Formal analysis:** Ya-Wei Eileen Lin, Tal Shnitzer, Ronen Talmon, Yuval Kluger.

**Funding acquisition:** Ronen Talmon, Kurt Schalper, Yuval Kluger.

**Investigation:** Ya-Wei Eileen Lin, Tal Shnitzer, Ronen Talmon, Kurt Schalper, Yuval Kluger.

**Methodology:** Ya-Wei Eileen Lin, Tal Shnitzer, Ronen Talmon, Kurt Schalper, Yuval Kluger.

**Project administration:** Ronen Talmon, Yuval Kluger.

**Resources:** Franz Villarroel-Espindola, Shruti Desai.

**Software:** Ya-Wei Eileen Lin, Tal Shnitzer.

**Supervision:** Ronen Talmon, Kurt Schalper, Yuval Kluger.

**Validation:** Ya-Wei Eileen Lin, Tal Shnitzer, Ronen Talmon, Kurt Schalper, Yuval Kluger.

**Visualization:** Ya-Wei Eileen Lin, Tal Shnitzer, Ronen Talmon, Yuval Kluger.

**Writing – original draft:** Ya-Wei Eileen Lin, Tal Shnitzer, Ronen Talmon, Yuval Kluger.

**Writing – review** & **editing:** Ya-Wei Eileen Lin, Tal Shnitzer, Ronen Talmon, Kurt Schalper, Yuval Kluger.

## Acknowledgments

The work of Y.-W E. Lin, T. Shnitzer and R. Talmon was supported by the European Union’s Horizon 2020 research grant agreement 802735.

The work of F. Villarroel-Espindola, S. Desai and K. Schalper was supported by was supported by the NIH grant R37CA245154, Yale SPORE in Lung Cancer P50CA196530, Stand Up To Cancer – American Cancer Society Lung Cancer Dream Team Translational Research Grants SU2C-AACR-DT1715 and SU2C-AACR-DT22-17, Department of Defense-Lung Cancer Research Program Career Development Award W81XWH-16-1-0160, Yale Cancer Center Support Grant P30CA016359, sponsored research by Navigate Biopharma and AstraZeneca.

The work of Y. Kluger was supported by NIH grant R01RGM131642, UM1DA051410 and P50CA121974.

The funders had no role in study design, data collection and analysis, decision to publish, or preparation of the manuscript.

## S1 Appendix

Given *S* is the set of the measurement. Based on the Taylor expansion, the kernel density estimation (KDE) for the sample *x*_*i*_:

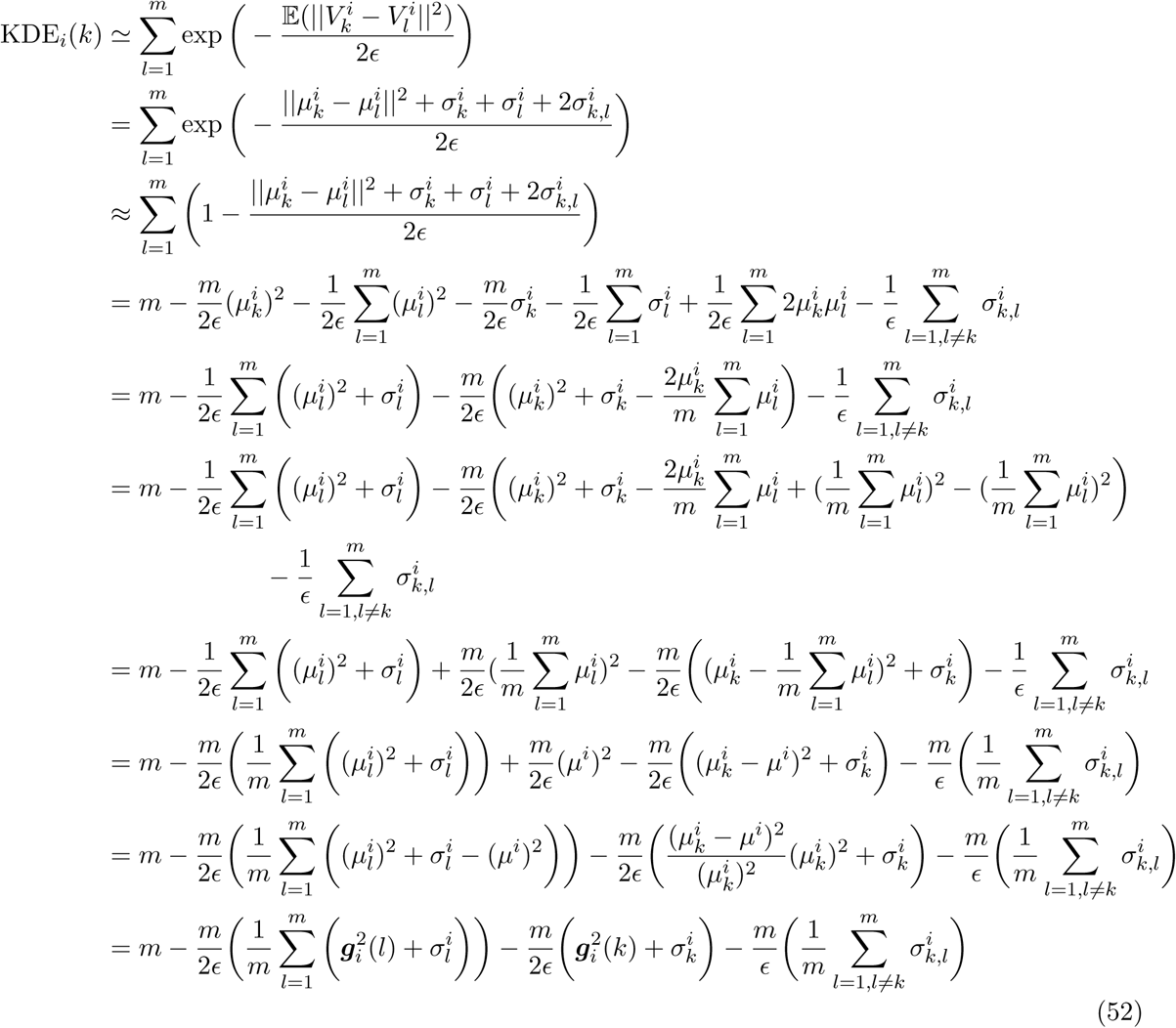

Note that ***π***_*i*_(*k*) *∝* ***D***_*i*_**1**, that is,

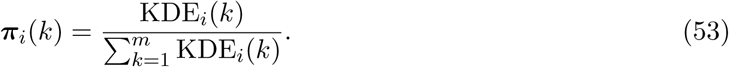

Therefore,

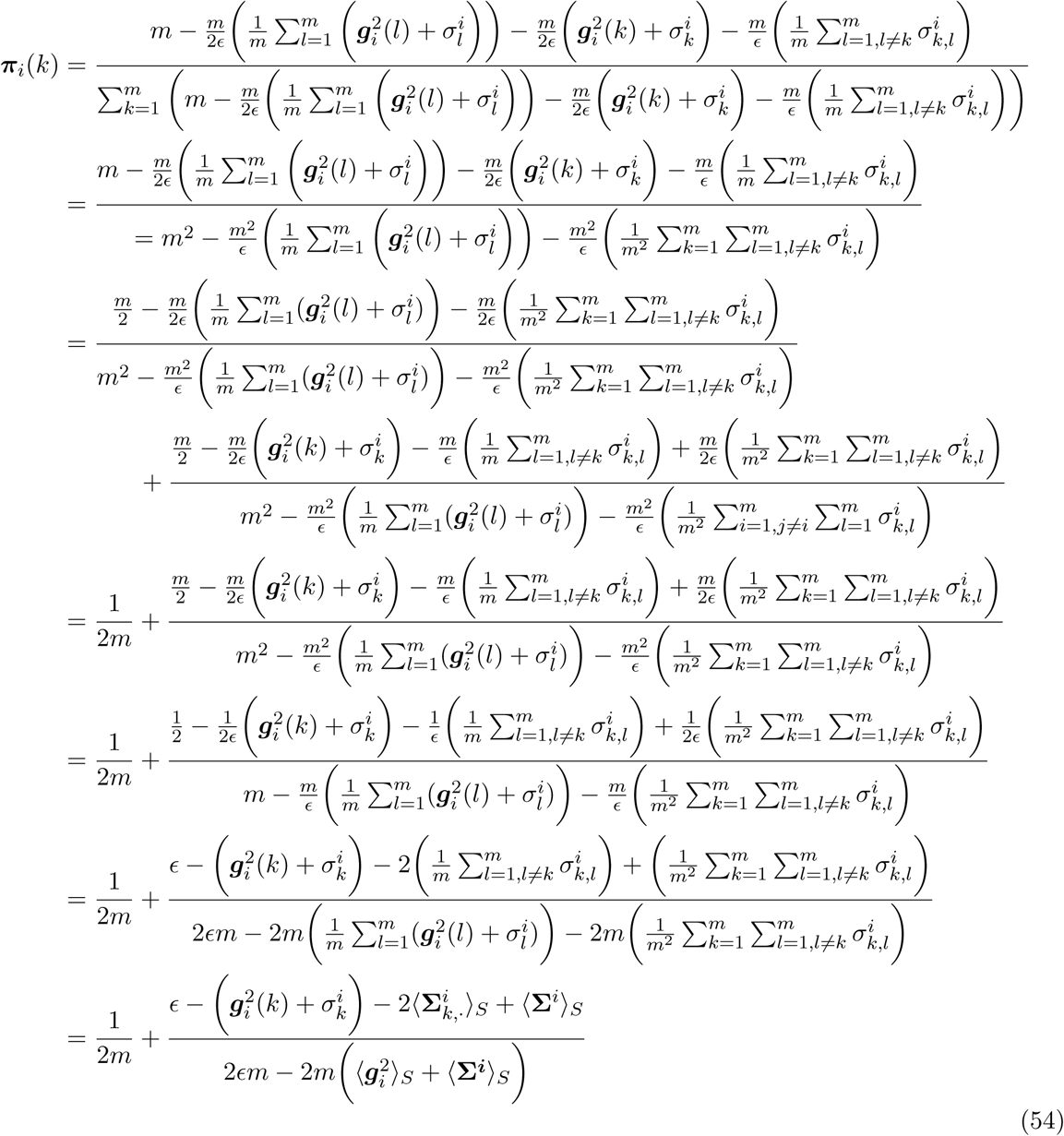

## Footnotes

1 The 29 markers are denoted by LipoR, VIM, T-BET, CD47, Cytokeratin, CD45RO, PD-L1, GAPDH, B7-H3, LAG-3, TIM-3, FOXP3, CD4, B7-H4, CD68, PD1, CD20, CD8, CD25, VISTA, KI-67, B2M, CD3, IDO-1, PD-L2, GZB, Histone 3, DNA1 and DNA2

